# Carbonic anhydrase 4 disruption and pharmacological inhibition reduce synaptic and behavioral adaptations following oxycodone withdrawal

**DOI:** 10.1101/2025.01.23.634619

**Authors:** Subhash C. Gupta, Rebecca J. Taugher-Hebl, Ali Ghobbeh, Marshal T. Jahnke, Rong Fan, Ryan T. LaLumiere, John A. Wemmie

## Abstract

The ongoing opioid crisis underscores the need for innovative treatments targeting the neurobiological mechanisms underlying opioid-seeking behaviors and relapse. Here, we explored the role of carbonic anhydrase 4 (CA4) in modulating synaptic adaptations to oxycodone withdrawal in mice. We disrupted CA4 genetically and inhibited it pharmacologically with acetazolamide (AZD), a carbonic anhydrase inhibitor used clinically. We found that oxycodone withdrawal increased AMPAR/NMDAR ratio and synaptic recruitment of calcium-permeable AMPARs in nucleus accumbens core (NAcC) medium spiny neurons (MSNs). Synaptic changes required an extended period of abstinence, generalized to other opioids, including morphine and heroin, were more pronounced in D1 dopamine receptor-expressing MSNs, and were prevented by CA4 disruption. AZD administration *in vitro* and *in vivo* reversed the synaptic alterations, and the effects of AZD depended on CA4 and acid-sensing ion channel-1A. Interestingly, abstinence from oxycodone did not affect dendritic spine density in NAcC MSNs, in contrast to previously observed effects of abstinence from cocaine. Finally, in an oxycodone self-administration paradigm, CA4 disruption and AZD reduced drug-seeking behaviors following 30 days of forced abstinence. Together, these findings identify a critical role for CA4 in synaptic adaptations in opioid withdrawn mice and drug-seeking behavior. Moreover, they suggest pharmacological inhibitors of CA4 may hold therapeutic potential for reducing opioid-seeking and relapse in opioid use disorder.

## Introduction

The opioid crisis remains a leading public health challenge, and overdose deaths still approach 100,000 per year. (https://www.cdc.gov/nchs/nvss/vsrr/drug-overdose-data.htm). Opioids and their withdrawal produce a sustained and increasing desire for more drugs, often referred to as opioid-seeking or craving [1]. Only a few medications are FDA-approved to treat opioid use disorder (OUD), and all of them target the mu opioid receptor [2]. Importantly, these medications do not correct the biological mechanisms that cause drug seeking or craving. Thus, new treatments with non-opioidergic mechanisms of action could have a substantial impact on the clinical management of OUD.

The nucleus accumbens core (NAcC) is a central hub in the brain circuits underlying responses to drugs of abuse and is thought to play a key role in the development of OUD [3, 4]. The majority of neurons (∼95%) in NAcC are medium spiny neurons (MSNs) [5, 6], which are critical for drug-seeking behaviors [7, 8]. Drugs of abuse, including cocaine and opioids, have been found to produce changes in synapses onto NAcC MSNs that persist well beyond acute drug exposures and may thus bias neural circuits towards drug seeking in the future, thus likely promoting vulnerability to relapse [9−12]. For example, cocaine withdrawal increases AMPAR/NMDAR ratio and Ca^2+^-permeable AMPARs (CP-AMPARs) at synapses onto NAcC-MSNs [10, 13−21], which is thought to promote cocaine-seeking behavior [7, 22]. Synaptic responses to opioids are less well-characterized, although some rearrangements similar to those evoked by cocaine have been reported. For example, withdrawal from non-contingent morphine administration increased AMPAR/NMDAR ratio in D1^+^ MSNs in NAc shell [23], and withdrawal from non-contingent administration of morphine, heroin, and oxycodone increased AMPAR/NMDAR ratio in NAcC-MSNs [24]. Supporting an important role for synaptic AMPARs in NAcC in promoting opioid-seeking behaviors, AMPAR antagonists delivered to the NAcC reduced reinstatement in rats following heroin self-administration [25].

We recently identified acid-sensing ion channels (ASICs) as a novel molecular mechanism in cocaine and opioid-seeking behaviors [20, 24]. ASICs are cation channels activated by extracellular acidosis that consist of trimeric assemblies of differing combinations of ASIC1A, ASIC2A, and ASIC2B subunits [26]. The ASIC1A subunit is required for activation by pH changes within a physiologically relevant range (below pH 7.4 to pH 5) [26]. In NAcC MSNs, we found that these channels are activated by protons released from glutamate-containing presynaptic vesicles during synaptic transmission [27, 28]. Loss of ASIC1A increased sensitivity of NAcC synapses to rearrangements induced by cocaine and opioids, and also increased conditioned place preference (CPP) to cocaine and opioids [18, 20, 24]. Together, these findings of heightened drug vulnerability in mice lacking ASIC1A suggest the intriguing possibility that increasing ASIC function might oppose synaptic and behavioral responses to both cocaine and opioids. Consistent with this possibility, overexpressing ASIC1A in NAcC in rats reduced cocaine self-administration. Subsequent work, however, suggested that overexpressing ASIC1A in NAcC in rats following cocaine self-administration and withdrawal increased reinstatement of cocaine-seeking behavior [29]. Thus, behavioral effects of potentiating ASIC function may be complex and depend on timing and neuron specificity.

Another potential strategy for increasing ASIC activation is to reduce pH buffering. Extracellular pH is buffered by the reaction (H^+^ + HCO_3_^−^↔CO_2_ + H_2_O) catalyzed by carbonic anhydrases (CA). Among more than 14 in mammals [30, 31], we focused on CA4 as a candidate for regulating synaptic ASICs based on several of its established characteristics. CA4 is abundantly expressed in brain neurons, including NAc MSNs, is anchored in the cell membrane facing the extracellular compartment, and has been previously suggested to buffer synaptic pH [32, 33]. Indeed, disrupting CA4 in postsynaptic MSNs increased ASIC-mediated EPSCs [18, 26]. Furthermore, loss of CA4 prevented synaptic rearrangements following cocaine withdrawal, including the associated increase in AMPAR/NMDAR ratio, CP-AMPARs, mEPSC frequency, and dendritic spine density [18, 20, 26]. Consistent with these synaptic effects of CA4 disruption, there were also behavioral effects. *Car4^−/−^* mice self-administered a similar amount of cocaine as *Car4^+/+^* mice, though after 4 weeks of forced abstinence, *Car4^−/−^* mice had reduced active lever presses (unreinforced by drug) [18]. CA4 disruption also reduced locomotor responses to acute cocaine challenge following cocaine withdrawal [18].

In this study, we investigated the effects of CA4 disruption on opioid withdrawal-induced synaptic adaptations and dendritic spine morphology in NAcC-MSNs, as well as oxycodone-seeking behavior. We also tested pharmacological inhibition of CA4 with acetazolamide (AZD), a carbonic anhydrase inhibitor used clinically for a variety of illnesses [34−36]. We hypothesized that CA4 disruption and pharmacological inhibition would protect against the effects of oxycodone withdrawal on glutamatergic synapses in NAcC and drug-seeking behavior. Our results suggest that CA4 may offer a new target for mitigating drug seeking and relapse in OUD that is unlike traditional opioid replacement therapies that target the mu opioid receptor.

## Materials and Methods

### Mice

All mice were on a C57BL/6 genetic background. *Car4*^−/−^ mice (stock no. 008217) [32] were obtained from The Jackson Laboratory. *Asic1a*^−/−^ mice were generated as previously described [37]. *Drd1a-tdTomato* mice (stock #016204) were sourced from Jackson Laboratory. Mice were housed in groups of 2−5 littermates with free access to standard chow and water. Mice were maintained on a 12-hour light-dark cycle with experiments performed during the light phase. Experimental groups were matched by sex and age (10−15 weeks). Within each housing group, mice were randomly assigned to treatment conditions. All mice were experimentally naïve at the start of each experiment. All animal procedures were approved by the University of Iowa Animal Care and Use Committee and complied with NIH guidelines for animal care.

### Oxycodone, heroin, morphine and AZD

Oxycodone, heroin, and morphine were kindly supplied by the National Institute on Drug Abuse. Acetazolamide (XGen, USA) was purchased from the University of Iowa Hospital pharmacy. The timing, dosing, and delivery of these drugs are described in the figures.

### Slice preparation and electrophysiology

Acute brain slices were prepared, and electrophysiological recordings were conducted following previously established protocols [18, 20, 24, 38]. AMPAR/NMDAR ratio was measured by recording evoked EPSCs, with the AMPAR-EPSC amplitude measured at −70 mV and NMDAR-EPSC amplitude at +50 mV, using the late component, 60 ms after onset. AMPAR rectification was evaluated by recording AMPAR-EPSCs at −70, −30, +30, and +50 mV, and plotting the current-voltage relationship. AMPAR rectification index was calculated as the ratio of current at −70 mV to current at +50 mV. To assess (N-((2-aminoethyl)-N, N-dimethyl) phenylpropan-2-amine) (NASPM) sensitivity, cells were held at −70 mV, and 20−25 baseline sweeps were collected. NASPM (200 μM, Alomone Lab) was applied, and 15 minutes later, 20−25 sweeps were collected. *In vitro* effects of CA4 inhibition were tested by applying AZD (100 μM) to the recording chamber for 1 hr.

### DiI labeling, dendritic spine imaging, and analysis

Mice were perfused, coronal slices cut, and NAcC MSNs were stained with DiI, as previously described [18, 38]. Dendritic segments were imaged, and spine density and morphology were analyzed using Neuron Studio, as previously described [18, 38]. Each experimental group included 3-4 animals, with 3-4 neurons analyzed per animal and 2-4 dendritic segments per neuron. Total number of spines assessed by group: *Car4*^+/+^ saline-veh (3850), *Car4*^+/+^ saline-AZD (4646), *Car4*^+/+^ oxycodone-veh (3007), and *Car4*^+/+^ oxycodone-AZD (3960).

### Oxycodone Conditioned Place Preference (CPP)

Oxycodone CPP was performed as previously described [24]. In brief, mice underwent pre- and post-tests during which they were allowed to explore both compartments of a two-chamber CPP apparatus (Med Associates) for 20 minutes. For 3 days between pre- and post-tests, mice underwent two training sessions per day during which oxycodone (15 mg/kg, i.p.) and saline were paired with opposite sides of the apparatus. Change in time spent on the oxycodone-paired side was calculated by subtracting the pre-test from the post-test.

### Oxycodone Self-Administration

To enable oxycodone self-administration, mice underwent jugular vein catheterization [18]. Catheter patency was verified post-surgery and during the course of the self-administration protocol. Mice were fasted overnight prior to self-administration training and were restricted to 85-90% of their pre-fasting body weight through sessions 1 to 10 to increase motivation [18]. Training occurred in operant conditioning chambers (Med Associates) equipped with cues (light and tone) and two levers (one active, one inactive). Mice self-administered oxycodone (0.25 mg/kg/infusion) on a fixed-ratio 1 (FR1) schedule during daily 6-hour sessions for a minimum of 10 days, based on a previously described protocol [39]. Active lever presses triggered an oxycodone infusion and 20 seconds of cues during which no additional infusions could be obtained. Mice meeting criteria of at least 10 infusions per day for 8 days, including the last 3 days, were included in further testing. 24 hours after the final self-administration session, baseline drug seeking was assessed by returning mice to the operant chambers for a 30-minute test session in which active lever presses triggered cues but no drug. After 30 days of forced abstinence, mice received an injection of AZD (30 mg/kg, i.p.) or vehicle (saline). 24 hours later, drug-seeking behavior was again assessed by returning mice to the operant chambers for a 30-minute test session in which active lever presses triggered cues but no drug. Active lever presses following abstinence were normalized to baseline to quantify change in drug-seeking behavior.

### Statistical analyses

Student’s t-test was used to assess statistical significance for experiments involving two groups. Two-way ANOVA was used to assess statistical significance for experiments involving 2 independent factors. ROUT test with Q = 1% was used to screen for outliers. The F-test was used to compare variances between groups, and Welch’s correction was applied for unequal variances. P values less than 0.05 were considered significant. All bar graphs express values as mean ± S.E.M. All statistical analyses were performed using GraphPad Prism.

## Results

To test whether CA4 disruption opposes synaptic responses to opioids, we delivered oxycodone (3 mg/kg, i.p.) vs. saline control once per day for 5 consecutive days to *Car4*^+/+^ and *Car4*^−/−^ mice and then withheld oxycodone for 10 days (**Fig. 1A**). We then harvested brain tissue and tested glutamatergic transmission onto NAcC-MSNs in acute brain slices (**Fig. 1B**). We found that oxycodone administration followed by abstinence increased AMPAR/NMDAR ratio in *Car4^+/+^* mice but not in *Car4*^−/−^ mice (**Fig. 1C-D**), suggesting that loss of CA4 prevented the oxycodone-induced change. These effects of oxycodone and CA4 disruption were independent of sex (**Fig. S1**). To test if extended abstinence was required to increase AMPAR/NMDAR ratio, we tested 24 hrs of abstinence after 5 doses (once per day for 5 days) and after one dose (**Fig. S2A**). The 24-hr abstinence periods after the 5 daily doses and after the single dose were not sufficient to change AMPAR/NMDAR ratio (**Fig. S2B**), suggesting an extended abstinence period was required.

**Figure 1:**
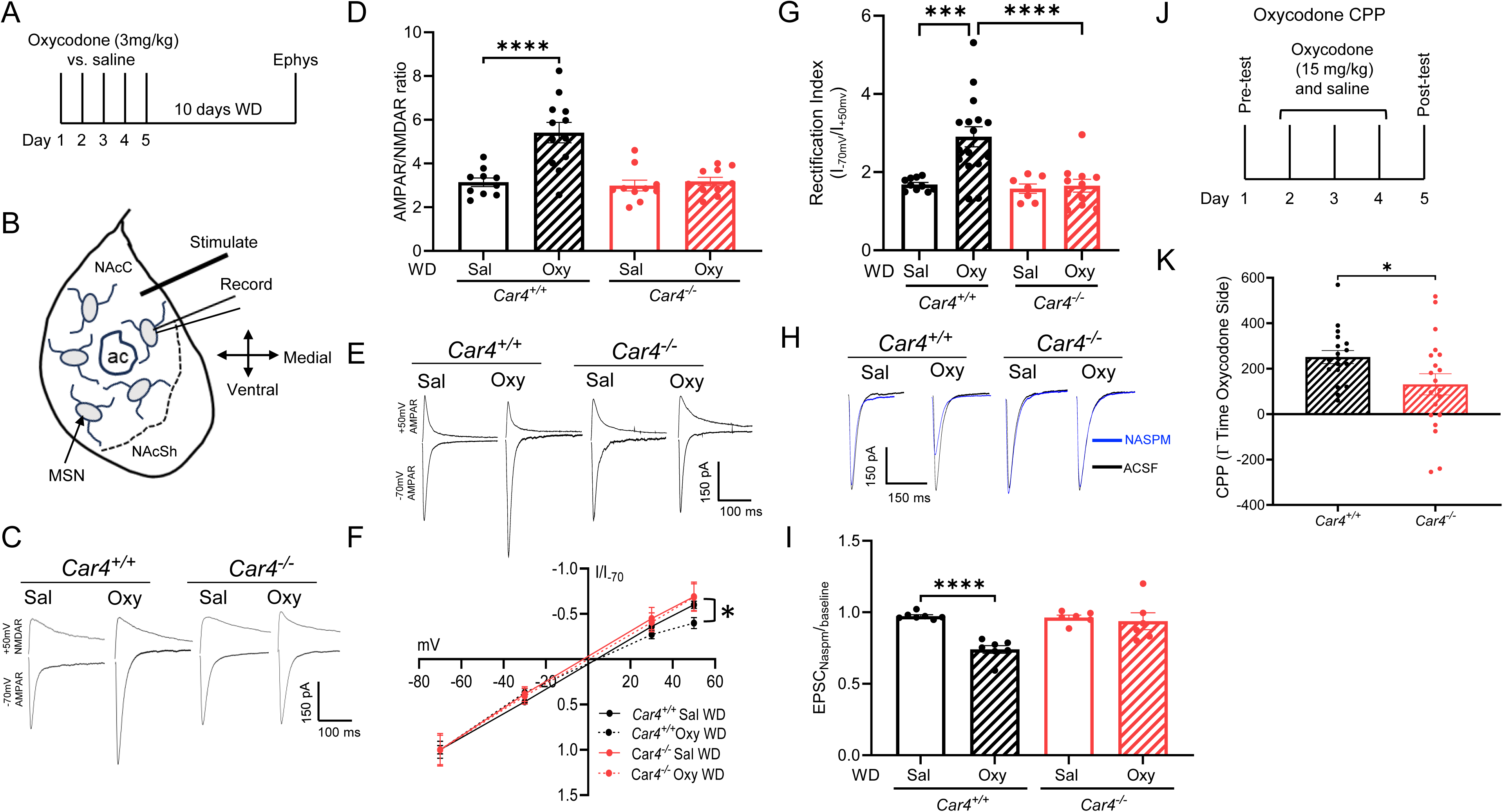
CA4 disruption protects against oxycodone withdrawal-induced synaptic changes at glutamatergic synapses in NAcC MSNs and reduces oxycodone conditioned place preference (CPP). (A) Experimental timeline: oxycodone (3mg/kg, i.p.) or saline (i.p.) was administered in the home cage each day for 5 days followed by 10 days of withdrawal, after which slice electrophysiology was performed. (B) Diagram illustrating the location of MSN recordings in NAcC and electrical stimulation. ac: anterior commissure; NAcSh: NAc shell. (C) Representative traces of the AMPAR-mediated EPSC at -70 mV and the NMDAR-mediated EPSC at +50 mV from NAcC MSNs from *Car4^+/+^* and *Car4^-/-^* mice following withdrawal from oxycodone (Oxy) vs saline (Sal). (D) AMPAR/NMDAR ratio increased after oxycodone withdrawal (Oxy) in *Car4^+/+^* but not in *Car4^-/-^* (2-Way ANOVA, drug by genotype interaction F(1, 38) = 10.55, p = 0.0024). Planned contrasts: Oxy vs. Sal *Car4^+/+^* ****p < 0.0001, Oxy vs. Sal *Car4^-/-^* p = 0.685. n = 10-12 neurons from 5 mice/group. (E) Example traces of AMPAR-mediated EPSCs at -70 mV and +50 mV in NAcC MSNs of *Car4^+/+^* and *Car4^-/-^* mice following oxycodone withdrawal (Oxy) vs saline withdrawal (Sal). (F) Current-voltage relationship (IV-curve): Oxycodone abstinence produced inward rectification in *Car4^+/+^* mice (drug by genotype interaction @ +50 mV F(1, 40) = 0.8833 p = 0.3529, drug F(1, 40) = 1.0 p = 0.3232, genotype F(1, 40) = 3.207 p = 0.0809). Planned contrasts: oxycodone withdrawal (Oxy WD) *Car4^+/+^* vs *Car4^-/-^*\*p = 0.0341. n = 7 to 16 neurons from 3-5 mice/group. (G) Quantification of rectification index (drug by genotype interaction F(1, 40) = 6.625, p = 0.0139). Planned contrasts: Oxy vs Sal *Car4^+/+^* ***p = 0.0001, Oxy vs Sal *Car4^-/-^* p = 0.8286, Oxy *Car4^+/+^* vs Oxy *Car4^-/-^* ****p < 0.0001. n = 7 to 16 neurons from 3 to 5 mice/group. (H) Representative traces of the AMPAR-mediated evoked EPSC at -70 mV before (black) and after (blue) NASPM treatment. (I) NASPM sensitivity increased after oxycodone withdrawal in *Car4^+/+^* but not in *Car4^-/-^* mice (drug by genotype interaction F(1, 22) = 10.40, p = 0.0039). Planned contrasts: Oxy vs Sal *Car4^+/+^* ****p < 0.0001, Oxy vs Sal *Car4^-/-^*p = 0.610. n = 6 to 7 neurons from 3 mice/group. (J) Experimental timeline for oxycodone CPP in *Car4^+/+^* and *Car4^-/-^* mice. (K) Oxycodone conditioned place preference is impaired in *Car4^-/-^* mice. *Car4^+/+^* vs *Car4^-/-^* (t(31.31) = 2.197, *p = 0.0355, n = 18, 20 mice/grp).

An increase in GluA2-lacking CP-AMPARs in the postsynaptic membrane is one mechanism that may contribute to an increase in AMPAR/NMDAR ratio [15, 40]. CP-AMPARs are more inwardly rectifying [13, 41]; therefore, we tested rectification index in NAcC MSNs by measuring current-voltage (I-V) relationships of synaptic AMPAR responses (**Fig. 1E**). Indeed, oxycodone abstinence increased rectification index in *Car4*^+/+^ mice relative to saline treated controls but not in *Car4*^−/−^ mice (**Fig. 1E-G**). To confirm these results, we also tested sensitivity to the CP-AMPAR-specific antagonist NASPM and found a significant increase in NASPM sensitivity in *Car4*^+/+^ mice following oxycodone withdrawal (**Fig. 1H, I**). Consistent with a protective effect of CA4 disruption, NASPM sensitivity was unchanged in *Car4*^−/−^ mice. Together, these findings suggest that oxycodone withdrawal increases CP-AMPARs in the postsynaptic membrane of NAcC MSNs and that loss of CA4 prevents this increase. These findings highlight CA4 disruption as a potential strategy for preventing synaptic rearrangements associated with oxycodone withdrawal.

To explore whether these effects of CA4 disruption might impact drug-seeking behaviors, we next tested oxycodone CPP, as previously described [24] (**Fig. 1J**). We found that both *Car4*^+/+^ and *Car4^−/−^* mice preferred the oxycodone-paired chamber, however, this preference was reduced in *Car4^−/−^* mice, suggesting that CA4 disruption reduced drug seeking (**Fig. 1 K**).

To explore whether pharmacologically inhibiting CA4 might produce effects similar to CA4 disruption, we focused on one of the best-characterized CA inhibitors, AZD [31, 42]. AZD is approved for the treatment of a variety of illnesses in humans [34−36, 43] and it has several properties that make it ideal for these studies. First, AZD potently blocks CA4 (∼10 nM affinity), as well as other CAs [44, 45]. Second, AZD readily crosses the blood-brain barrier [46]. Moreover, it is rapidly metabolized and excreted by the kidneys, with a half-life of approximately 1 hr in mice [47, 48]. Finally, AZD potentiates ASIC1A-mediated synaptic currents in NAcC MSNs in a CA4-dependent manner [18, 20].

To determine whether AZD might produce effects similar to CA4 disruption, we first tested its effects in brain slices in *Car4^+/+^* mice. We delivered oxycodone (3 mg/kg, i.p. vs. saline) daily for 5 days, followed by 5 days of abstinence (**Fig. 2A**). We then assessed AMPAR/NMDAR ratio in NAcC MSNs. As before, oxycodone withdrawal increased AMPAR/NMDAR ratio (**Fig. 2B, C**). Compared to the vehicle, applying AZD (100 μM) to the bath for 1 hr significantly reduced AMPAR/NMDAR ratio in oxycodone withdrawn mice (**Fig. 2B, C**). Importantly, AZD had no effect on AMPAR/NMDAR ratio in saline-treated mice. To test generalizability to other opioids, we delivered morphine (10 mg/kg, i.p. vs. saline) daily for 10 days, followed by 5 days of abstinence and again applied AZD in the bath (**Fig. 2D**). AZD treatment similarly protected against the increase in AMPAR/NMDAR ratio induced by abstinence from morphine (**Fig. 2D-F**). Together, these data suggest that within 1 hr, acute application of AZD to brain slices can normalize opioid-induced increases in AMPAR/NMDAR ratio.

**Figure 2:**
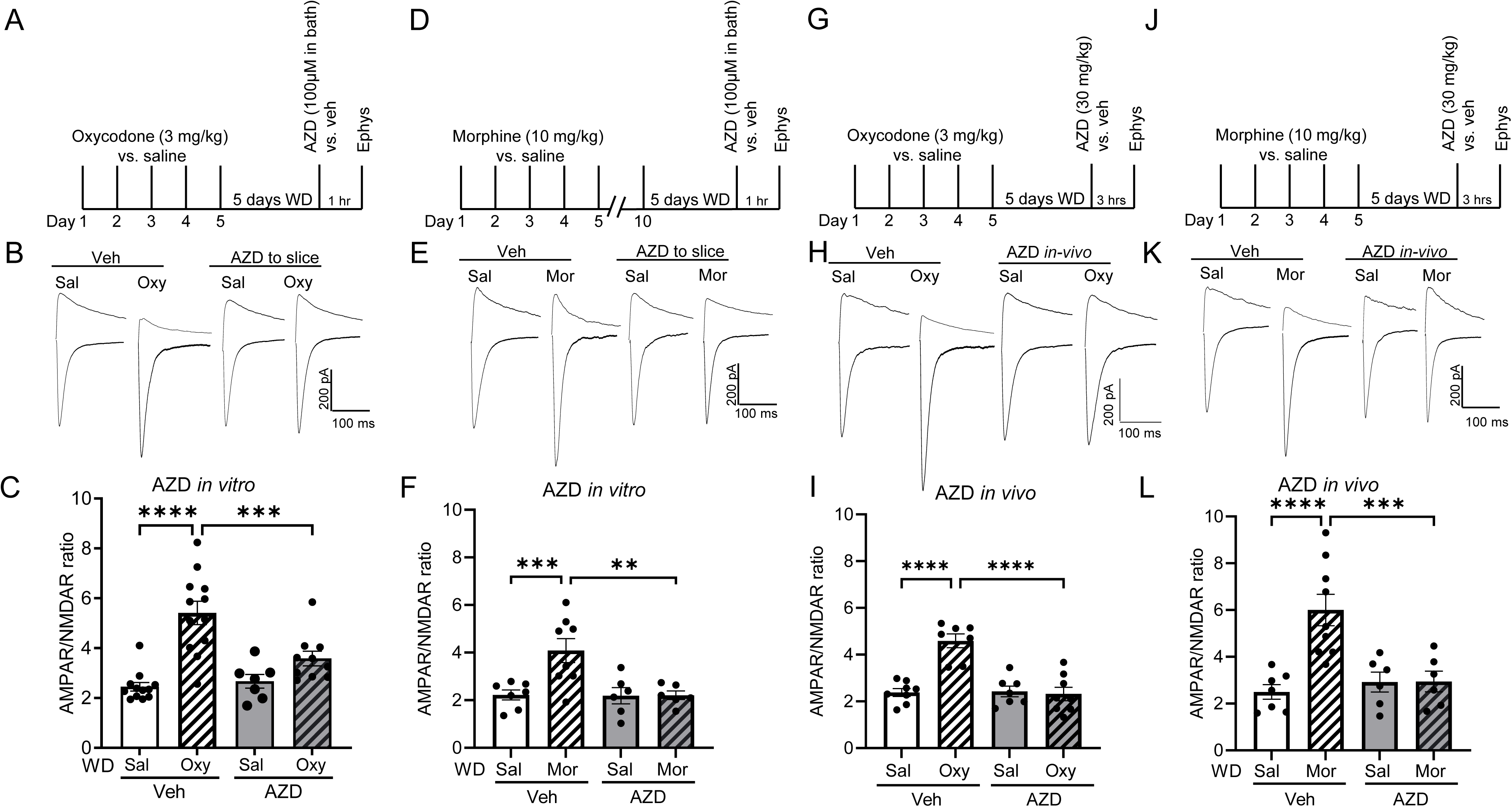
*In vitro* and *in vivo* administration of the CA inhibitor AZD reversed effects of opioid abstinence on AMPAR/NMDAR ratio. (A) Experimental timeline: oxycodone (3mg/kg, i.p.) or saline (i.p.) was administered each day for 5 days, followed by 5 days of withdrawal. Brain slices were prepared, and AZD 100µM or vehicle (ACSF) was added to the recording chamber for 60 minutes before testing. (B) Representative traces of AMPAR/NMDAR from *Car4^+/+^* mice after saline (Sal) and oxycodone withdrawal (Oxy) and in vitro treatment with AZD or vehicle (ACSF). (C) The AMPAR/NMDAR ratio increased after oxycodone withdrawal (Oxy) in *Car4^+/+^* mice. Applying AZD to the recording chamber reversed values to control levels (Oxy by AZD interaction F(1, 37) = 8.729, p = 0.0054). Planned contrasts: Oxy-Veh vs Sal-Veh ****p < 0.0001, Oxy-Veh vs Oxy-AZD ***p = 0.0003, Oxy-Veh vs Sal-AZD ****p < 0.0001). n *=* 7-12 neurons from 3-4 mice/group. (D) Experimental timeline: morphine (10mg/kg) or saline (i.p.) was administered each day for 10 days, followed by 5 days of withdrawal. Brain slices were prepared, and AZD 100µM or vehicle (ACSF) was added to the recording chamber for 60 minutes before testing. (E) Representative traces of AMPAR/NMDAR from *Car4^+/+^* mice after saline (Sal) and morphine withdrawal (Mor) and *in vitro* treatment with AZD or vehicle (ACSF). (F) AMPAR/NMDAR ratio increased after morphine withdrawal (Mor) in *Car4^+/+^* mice. AZD reversed this measure to control levels (Mor by AZD interaction F(1, 23) = 6.291, p = 0.0196). Planned contrasts: Mor-Veh vs Sal-Veh ***p = 0.0009, Mor-Veh vs Mor-AZD **p = 0.0013, Mor-Veh vs Sal-AZD **p = 0.0012. n *=* 6-8 neurons from 3 mice/group. (G) Experimental timeline: oxycodone (3 mg/kg vs. saline (i.p, 5 days), followed by 5 days of withdrawal. 3 hrs later, AZD (30 mg/kg vs. vehicle was administered *in vivo*, and slices were harvested for electrophysiological recording. (H) Representative traces of AMPAR/NMDAR from *Car4^+/+^* mice after saline (Sal) and oxycodone withdrawal (Oxy) and following *in vivo* treatment with AZD vs. vehicle. (I) AZD treatment reversed the oxycodone withdrawal-induced increase in AMPAR/NMDAR ratio (Oxycodone by AZD interaction F(1, 26) = 21.69, p < 0.0001). Planned contrasts: Oxy-Veh vs. Sal-Veh ****p < 0.0001, Oxy-Veh vs Oxy-AZD ****p < 0.0001. n *=* 7-8 neurons from 3 mice/group. (J) Experimental timeline: morphine (10mg/kg), or saline (i.p.) was administered for 5 days, followed by 5 days of withdrawal. AZD (30/mg/kg) or vehicle was administered by i.p. injection in vivo, and 3 hrs later, slices were harvested for electrophysiological recording. (K) Representative traces of AMPAR/NMDAR from *Car4^+/+^* mice after saline (Sal) and morphine withdrawal (Mor) and following *in vivo* treatment with AZD vs. vehicle. (L) *In vivo* administration of AZD reversed AMPAR/NMDAR ratios following morphine abstinence to control levels (Mor by AZD interaction F(1, 24) = 10.34, p = 0.0196). Planned contrasts: Mor-Veh vs Sal-Veh ****p < 0.0001, Mor-Veh vs Mor-AZD ***p = 0.0004. n *=* 6-9 neurons from 3 mice/group.

We next tested whether administering AZD *in vivo* might have similar effects. We estimated that an AZD dose of 30 mg/kg, i.p. should achieve a systemic concentration similar to that used in our brain slice experiments and analogous to that safely used in humans [48]. We administered oxycodone (3 mg/kg, i.p. vs. saline) daily for 5 days. After 5 days of abstinence (**Fig. 2G**), we gave AZD vs. vehicle. Three hours later, we harvested brain slices and assessed AMPAR/NMDAR ratio in NAcC MSNs. Similar to our observations with AZD *in vitro*, administering AZD *in vivo* after oxycodone withdrawal reduced AMPAR/NMDAR to levels comparable to vehicle-treated, saline-withdrawn counterparts (**Fig. 2H, I**), AZD had no effect in mice not exposed to oxycodone. We also tested the effects of *in vivo* AZD administration after 5 days of withdrawal from morphine and observed similar results (**Fig. 2J-L**).

Next, we tested whether the effects of the *in vivo*-administered AZD lasted beyond 3 hrs. We administered AZD vs. vehicle following abstinence from oxycodone and tested AMPAR/NMDAR 24 hrs later (**Fig. 3A**). As before, we found that oxycodone withdrawal significantly increased AMPAR/NMDAR ratio in vehicle-treated *Car4*^+/+^ mice (**Fig. 3B**). Importantly, this change was absent in the AZD-treated mice (**Fig. 3B**), suggesting that AZD reversed AMPAR/NMDAR ratio back to baseline levels. AZD administration had no effect in controls receiving saline instead of oxycodone (**Fig. 3B**), suggesting that the effects of AZD were specific to synaptic changes induced by oxycodone withdrawal. Additionally, as with CA4 disruption, the effects of AZD were independent of sex **(Fig. S1)**. Moreover, these synaptic effects of AZD likely persisted after AZD was cleared because AZD’s half-life is short (∼ 1hr) [47, 48]. Together, these findings suggest that AZD opposes-opioid induced synaptic rearrangements resembling effects of genetically disrupting CA4.

**Figure 3:**
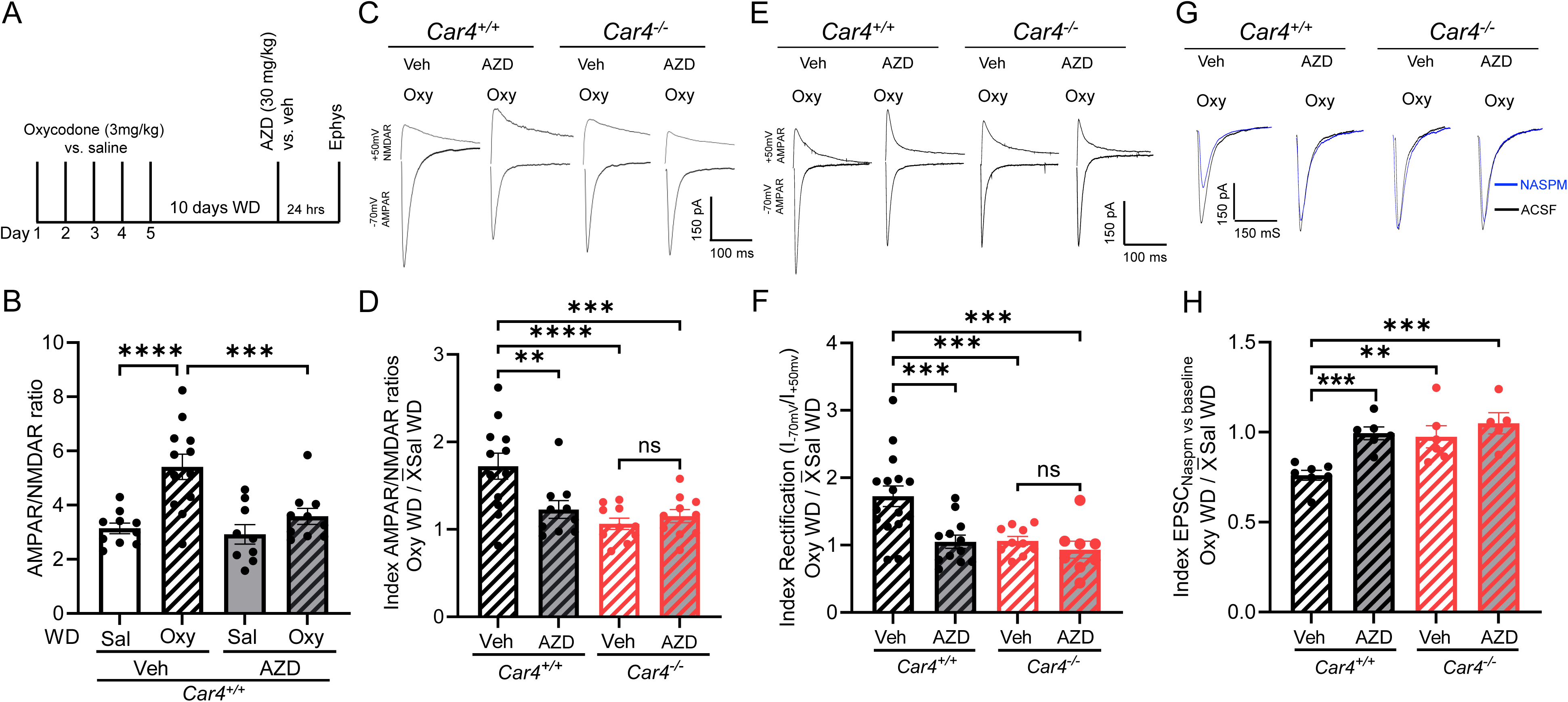
Acetazolamide (AZD) reversed oxycodone withdrawal-induced changes at glutamatergic synapses in NAcC MSNs in *Car4^+/+^* mice but not in *Car4^−/−^* mice. (A) Experimental timeline: oxycodone (3mg/kg, i.p.) or saline (i.p.) was administered each day for 5 days, followed by 10 days of withdrawal. AZD (30/mg/kg, i.p.) or vehicle (saline, i.p.) was administered, and 24-hrs later, slices were harvested for electrophysiological recording. (B) AZD treatment reversed AMPAR/NMDAR ratios in oxycodone-withdrawn mice to control levels in saline-withdrawn mice (Oxy by AZD interaction F(1, 37) = 4.966, p = 0.032). Planned contrasts: Oxy-Veh vs Sal-Veh ****p < 0.0001, Oxy-Veh vs Oxy-AZD ***p = 0.0007. n *=* 10-12 neurons from 4 mice/group. (C) Representative traces of AMPAR/NMDAR from *Car4^+/+^* and *Car4^-/-^* mice after withdrawal from oxycodone (Oxy) or saline (Sal) and treatment with AZD vs. vehicle (Veh). (D) AZD significantly reduced AMPAR/NMDAR ratio in *Car4^+/+^* mice but not in *Car4^-/-^* mice. Indices of AMPAR/NMDAR ratio in oxycodone-withdrawn mice divided by means of saline withdrawn counterparts are shown (Oxy by AZD interaction F(1, 38) = 7.189, p = 0.0108). Planned contrasts: *Car4^+/+^*, Oxy WD-Veh vs Oxy WD-AZD **p = 0.0021; *Car4^+/+^* Oxy WD-Veh vs *Car4^-/-^* Oxy WD-Veh ****p < 0.0001, *Car4^+/+^* Oxy WD-Veh vs Car4^-/-^ Oxy WD-AZD ***p = 0.0005. n = 10-12 neurons from 4 mice/group. (E) Representative traces of AMPAR-mediated EPSCs at -70 and +50 mV from oxycodone withdrawn *Car4^+/+^* and *Car4^-/-^* mice after treatment with AZD vs. vehicle. (F) AZD significantly reduced AMPAR rectification relative to vehicle treatment in *Car4^+/+^* but not in *Car4^-/-^* mice. Indices of rectification in oxycodone-withdrawn mice divided by means of saline withdrawn counterparts are shown (Oxy by AZD interaction F(1, 38) = 7.189, p = 0.0108). Planned contrasts: *Car4^+/+^*, Oxy WD-Veh vs Oxy WD-AZD ***p = 0.0003; *Car4^+/+^* Oxy WD-Veh vs *Car4^-/-^* Oxy WD-AZD ***p = 0.0006; *Car4^+/+^* Oxy WD-Veh vs *Car4^-/-^* Oxy WD-Veh ***p = 0.0002. n = 10-12 neurons from 4 mice/group. (G) Representative traces of the AMPAR-mediated evoked EPSC at -70 mV before (black) and after (blue) NASPM application from *Car4^+/+^* and *Car4^-/-^*mice after oxycodone withdrawal (Oxy) and treatment with AZD vs. vehicle. (H) AZD treatment normalized NASPM sensitivity in *Car4^+/+^* mice not in *Car4^-/-^* mice. Indices of EPSCs in presence of NASPM vs baseline in oxycodone-withdrawn mice divided by means of saline-withdrawn counterparts are shown (Oxy by AZD interaction F(1, 22) = 4.617, p = 0.0429). Planned contrasts: *Car4^+/+^*, Oxy WD-Veh vs Oxy WD-AZD ***p = 0.0003; *Car4^+/+^* Oxy WD-Veh vs *Car4^-/-^*Oxy WD-Veh **p = 0.0032; *Car4^+/+^* Oxy WD-Veh vs *Car4^-/-^*Oxy WD-AZD *\*\*\**p = 0.0002. n = 6-7 neurons from 3 mice/group.

We hypothesized that the effects of AZD are mediated through its ability to inhibit CA4 rather than another target. To test if AZD acts via CA4, we compared the effects of AZD in *Car4^+/+^* and *Car4^−/−^* mice. We found that while AZD significantly reduced AMPAR/NMDAR in oxycodone-withdrawn *Car4^+/+^* mice, it had no effect in *Car4^−/−^*mice (**Figs. 3C-D and S3A-B)**. These results suggest effects of AZD are mediated through CA4.

We similarly tested the effects of AZD on the oxycodone withdrawal-induced changes in AMPAR rectification and NASPM sensitivity. In *Car4*^−/−^ mice, no effects of oxycodone withdrawal or AZD were observed (**Figs. 3E-H** and **S3C-F**). This lack of AZD responses in *Car4*^−/−^ mice further supports our hypothesis that the effects of oxycodone and AZD depend on CA4.

To test whether the protective effects of AZD against withdrawal-induced synaptic adaptations generalized to other opioids, we also tested withdrawal from heroin (8mg/kg for 5 days) and morphine (10mg/kg for 5 days) (**Fig. 4A**). After 10 days of withdrawal, mice received AZD or vehicle (saline), and 24 hours later AMPAR/NMDAR ratio was assessed. Withdrawal from both heroin (**Fig. 4B, C**) and morphine (**Fig. 4D, E**) significantly increased AMPAR/NMDAR ratio in NAcC-MSNs. Moreover, AZD reversed these increases (**Fig. 4B-E**). Withdrawal from a lower heroin dose (2 mg/kg, i.p.) did not significantly increase AMPAR/NMDAR ratio (**Fig. S4A-4C**), suggesting that opioid-induced synaptic rearrangements are dose-dependent. Interestingly, the same synaptic rearrangements and AZD effects were also observed after just 5 days of withdrawal from morphine (**Fig. S5A-5C**), suggesting that abstinence periods as short as 5 days, but longer than 24 hrs, are sufficient to alter AMPAR/NMDAR ratio. Together, these observations suggest that synaptic adaptations evoked by oxycodone withdrawal and the normalizing effects of AZD are not specific to oxycodone but generalize to glutamatergic plasticity evoked by other opioids.

**Figure 4:**
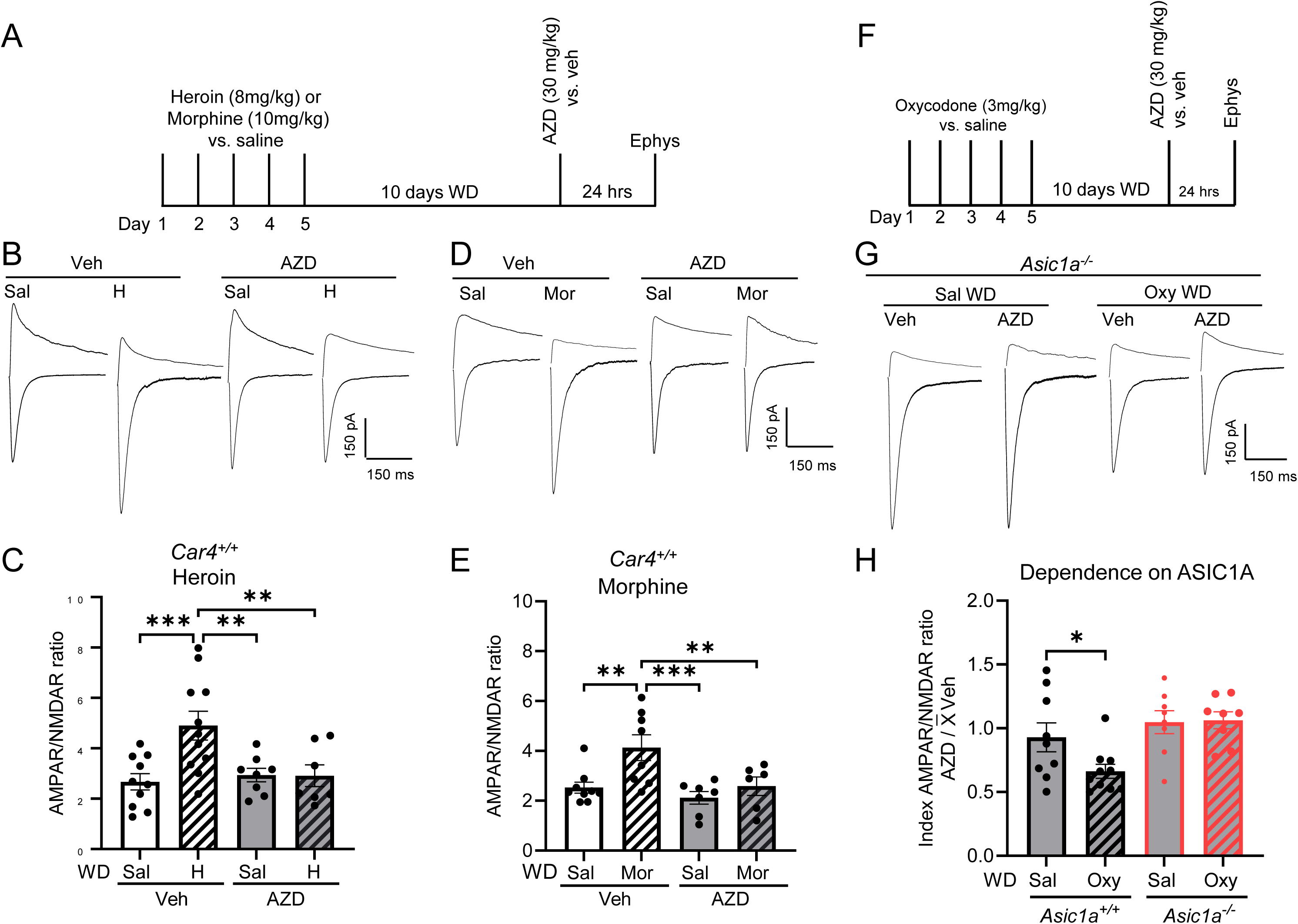
Effects of AZD on oxycodone-induced increases in AMPAR/NMDAR depend on ASIC1A and generalize to other opioids. (A) Experimental timeline for heroin, morphine, and AZD treatment. (B) Representative traces of AMPAR/NMDAR from *Car4^+/+^* mice after heroin (H) or saline (Sal) withdrawal and treatment with AZD vs vehicle. (C) AZD treatment reversed the heroin-induced increase in AMPAR/NMDAR ratio relative to vehicle-treated controls (H by AZD interaction F(1, 32) = 6.156, p = 0.0185). Planned contrasts: Sal-Veh vs H-Veh ***p = 0.0006, H-Veh vs H-AZD **p = 0.0043, and H-Veh vs Sal AZD **p = 0.0034. n = 8-11 neurons from 4 mice/group. (D) Representative traces of AMPAR/NMDAR from *Car4^+/+^* mice after morphine (Mor) or Saline (Sal) withdrawal (Mor) and treatment with AZD vs. vehicle (E) AZD treatment reversed morphine-induced increase in AMPAR/NMDAR ratio compared to vehicle-treated controls (Mor by AZD interaction F(1, 26) = 2.452, p = 0.1295; Mor F(1, 26) = 8.088, p = 0.0086; AZD F(1, 26) = 7.179, p = 0.0126). Planned contrasts: Sal-Veh vs Mor-Veh **p = 0.0025, Mor-Veh vs Mor-AZD **p = 0.0074, and Mor-Veh vs Sal-AZD ***p = 0.0005. n = 7-9 neurons from 4 mice/group. (F) Experimental timeline: oxycodone (3mg/kg, i.p.) or saline (i.p.) was administered each day for 5 days, followed by 10 days of withdrawal. AZD or vehicle was administered, and 24-hrs later, slices were harvested for electrophysiological recording. (G) Representative traces of AMPAR/NMDAR from *Asic1a^-/-^* mice withdrawn from oxycodone (Oxy) or saline (Sal) and treated with AZD vs. vehicle. (H) Index of AMPAR/NMDAR in AZD-treated mice, relative to the mean of vehicle-treated counterparts. This index of the AZD effect was reduced following oxycodone withdrawal in *Asic1a^+/+^*mice, but not *Asic1a^-/-^* mice. There was a significant effect of genotype (F(1, 31) = 9.485, p = 0.0043), but not oxycodone (F(1, 31) = 2.269, p = 0.1421) and no interaction (F(1, 31) = 2.828, p = 0.1027). Planned contrasts: *Asic1a^+/+^* Sal vs Oxy *p = 0.025, *Asic1a^-/-^* Sal vs Oxy p = 0.9059. n = 8-10 neurons from 4 mice/group.

To test whether the effects of AZD depended on ASIC1A, we administered oxycodone (3mg/kg) vs. saline to *Asic1a^+/+^* and *Asic1a^-/-^*mice for 5 days and then withheld it for 10 days (**Fig. 4F**). We then tested AMPAR/NMDAR ratio in mice treated with AZD vs. vehicle. In *Asic1a^-/-^* mice, AZD had no effect regardless of whether the mice underwent oxycodone withdrawal or not (**Figs. 4G, H** and **S6**). Together, these observations suggest that the ability of AZD to reduce AMPAR/NMDAR ratio following oxycodone withdrawal depends on both ASIC1A and CA4.

NAcC MSNs are subclassified by expression of different dopamine receptor subtypes (D1 vs. D2) [5, 6]. These different MSN subtypes have been suggested to play differing roles, with D1-expressing (D1^+^-NAcC-MSNs) promoting drug-seeking behaviors after withdrawal [7, 21, 49, 50], and D2-expressing opposing drug seeking [7, 8]. Therefore, we asked which subtype of NAcC-MSNs exhibited the glutamatergic changes following oxycodone withdrawal. To answer this question, we leveraged reporter mice expressing tdTomato specifically in D1^+^ MSNs (**Fig. 5A**). We tested responses in AMPAR/NMDAR ratio in D1^+^-NAcC-MSNs versus non-D1^+^-NAcC-MSNs (not expressing tdTomato) using the same oxycodone and AZD exposures as before (**Fig. 3A**). We performed recording from D1^+^, and non-D1^+^ NAcC-MSNs as illustrated (**Fig. 5B**). We found oxycodone withdrawal evoked an increase in AMPAR/NMDAR ratio in D1^+^, but not in non-D1^+^ NAcC-MSNs (**Fig. 5C-F**). Notably, AZD reversed the oxycodone evoked increase in AMPAR/NMDAR in D1^+^ MSNs and had no effect on non-D1^+^ MSNs (**Fig. 5C-F**).

**Figure 5.**
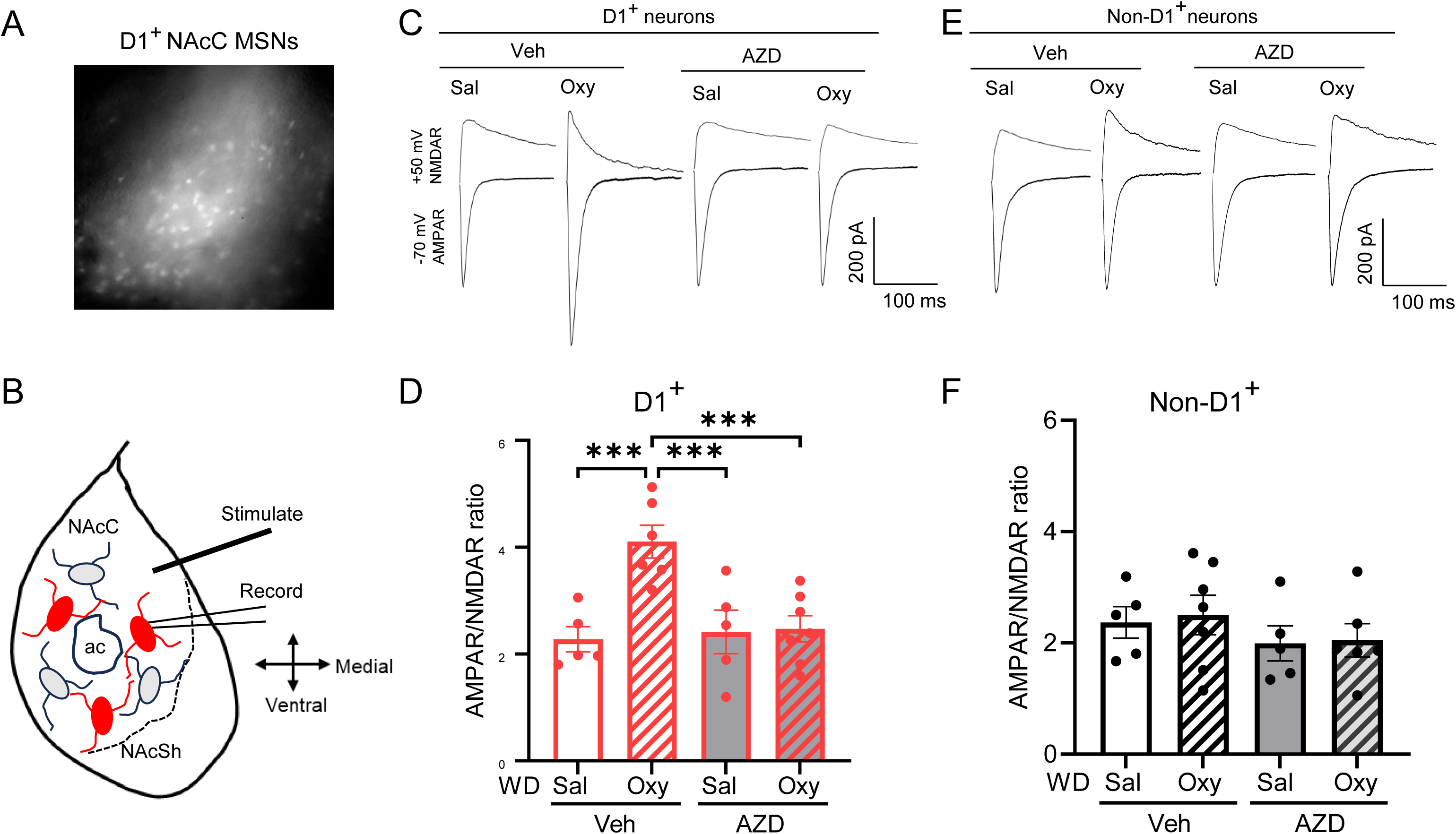
Increase in AMPAR/NMDAR ratio in oxycodone withdrawn mice was specific to D1^+^ neurons. (A) Image of D1^+^ neurons in NAcC labeled with tdTomato. (B) Diagram illustrating the location of recordings of D1^+^ and non-D1^+^ MSNs recording in NAcC. (C) Representative traces of AMPAR/NMDAR ratio of D1^+^ neurons in oxycodone withdrawn (Oxy) and saline withdrawn (Sal) mice treated with AZD vs. vehicle. (D) Oxycodone withdrawn mice exhibit increases in AMPAR/NMDAR ratio D1^+^ neurons, and AZD administration reversed the measure to levels in vehicle-treated controls (Oxy by AZD interaction F(1, 19) = 8.578 p = 0.0086). Planned contrasts: Oxy-Veh vs Sal-Veh ***p = 0.0005; Oxy-Veh vs Oxy-AZD ***p = 0.0006, and Oxy-Veh vs Sal-AZD **p = 0.001. n = 5-7 neurons from 3 mice/group. (E) Representative traces of AMPAR/NMDAR ratio of non-D1^+^ MSNs in oxycodone withdrawn (Oxy) and saline withdrawn (Sal) mice treated with AZD vs. vehicle. (F) Oxycodone withdrawn mice exhibited no change AMPAR/NMDAR ratio in D1^−^ neurons, and AZD had no effect. (Oxy by AZD interaction F(1, 19) = 0.157 p = 0.9017; Oxy F(1, 19) = 0.0838 p = 0.7754; AZD F(1, 19) = 1.609 p = 0.22). Planned contrasts: Oxy-Veh vs Sal-Veh p = 0.7688; Oxy-Veh vs Oxy-AZD p = 0.3047, and Oxy-Veh vs Sal-AZD p = 0.2765. n = 5-7 neurons from 3 mice/group.

Synaptic rearrangements following withdrawal from cocaine are accompanied by increases in dendritic spine density in NAcC MSNs [51−53] that may contribute to drug-seeking behaviors [53, 54]. Effects of opioid withdrawal on dendritic spines are less well characterized, although recent results suggest that the effects of opioids may differ from those of cocaine [24, 53]. Because CA4 disruption attenuated the increased dendritic spine density following cocaine withdrawal[18], we were interested in whether oxycodone withdrawal and inhibiting CA4 with AZD would affect dendritic spine density. To test this possibility, we administered oxycodone for 5 days, followed by 10 days of abstinence. We then administered AZD or vehicle and 24 hrs later harvested brain tissue for dendritic spine analyses (**Fig. 6A**). We found that total spine density was similar between oxycodone- and saline-withdrawn *Car4^+/+^*mice (**Fig. 6B, C**), with no differences in the density of stubby (**Fig. 6D**), thin (**Fig. 6E**), or mushroom spines (**Fig. 6F**). These results were largely consistent with the recently reported absence of effect of oxycodone withdrawal on dendritic spine density in NAcC-MSNs [24]. Importantly, in both oxycodone- and saline-withdrawn mice, AZD had no effect on any of these measures versus vehicle (**Fig. 6B-F**). Together, these results suggest that oxycodone withdrawal exerted little or no effect on dendritic spine density in NAcC-MSNs, and thus its effects are distinct from those of cocaine withdrawal. In addition, these data suggest that alterations in glutamatergic neurotransmission evoked in NAcC-MSNs by oxycodone withdrawal are not accompanied by effects on spine density.

**Figure 6.**
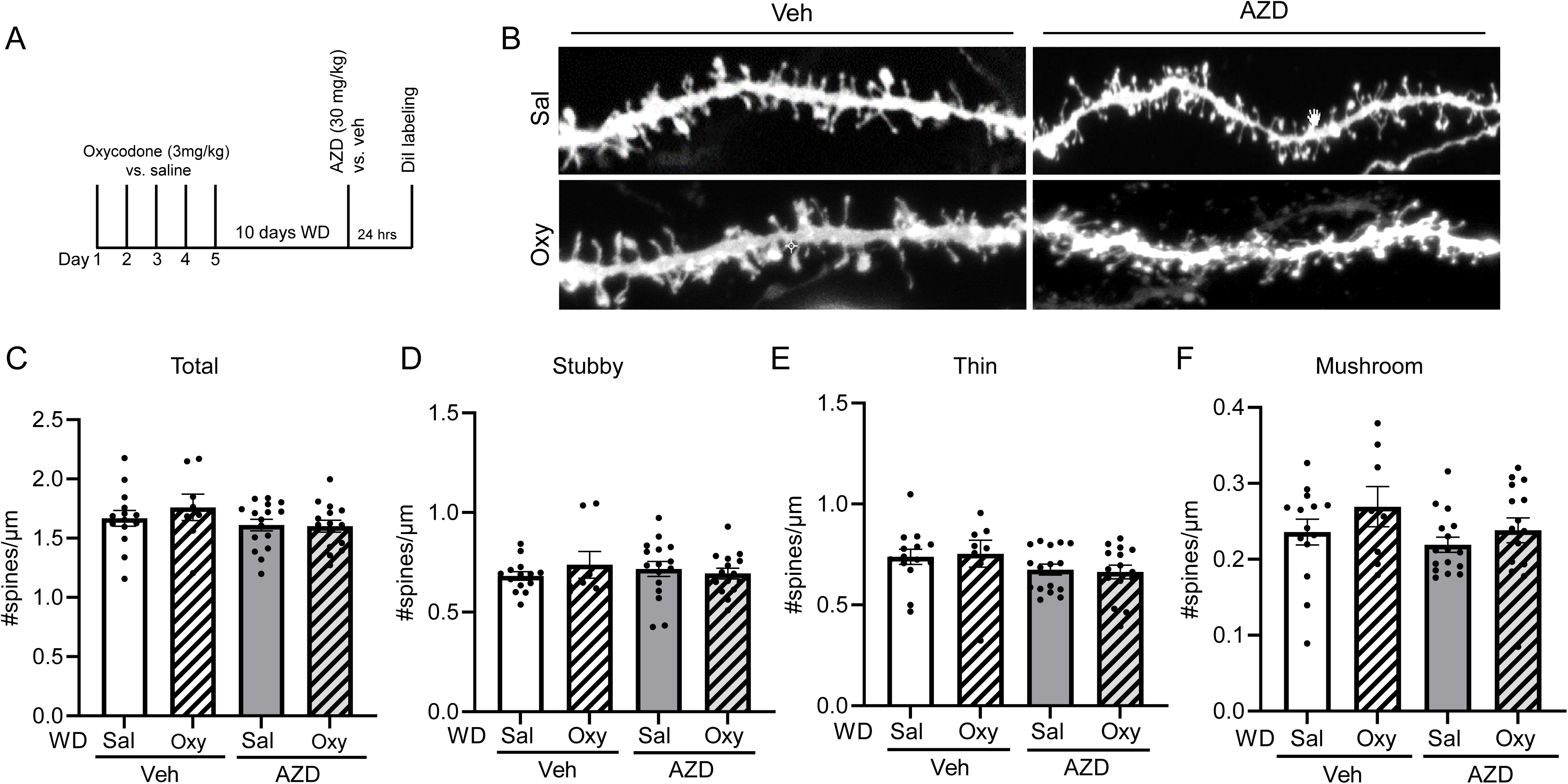
Oxycodone withdrawal and AZD had no effect on dendritic spine densities in NAcC MSNs. (A) Experimental timeline: oxycodone (3mg/kg, i.p.) or saline (i.p.) was administered in the home cage each day for 5 days, followed by 10 days of withdrawal, after which DiI labeling was performed. (B) Representative images (projected z-stack) of dendritic spines in NAcC MSNs from *Car4^+/+^* mice withdrawn from oxycodone (Oxy) vs saline (Sal) and treated with AZD vs. vehicle. (C) Total spine density was unchanged by oxycodone withdrawal and AZD (Oxy by AZD interaction F(1, 49) = 0.6042 p = 0.4407; Oxy F(1, 49) = 0.3975 p = 0.5313; AZD F(1, 49) = 2.694 p = 0.1071). Planned contrasts: Oxy-Veh vs Sal-Veh p = 0.3692, Oxy-Veh vs Oxy-AZD p = 0.1214; Sal-Veh vs Sal-AZD p = 0.5057. n = 8 to 16 neurons from 3 to 5 mice/group. (D) Stubby spine density was unchanged by oxycodone withdrawal and AZD (Oxy by AZD interaction F(1, 49) = 1.121 p = 0.2949; Oxy F(1, 49) = 0.2006 p = 0.6562; AZD F(1, 49) = 0.0138 p = 0.9071). Planned contrasts: Oxy-Veh vs Sal-Veh p = 0.3368, Oxy-Veh vs Oxy-AZD p = 0.4471; Sal-Veh vs Sal-AZD p = 0.4668. (E) Thin spine density was unchanged by oxycodone withdrawal and AZD (Oxy by AZD interaction F(1, 49) = 0.1144 p = 0.7367; Oxy F(1, 49) = 0.0033 p = 0.9542; AZD F(1, 49) = 3.799 p = 0.0570). Planned contrasts: Oxy-Veh vs Sal-Veh p = 0.7999, Oxy-Veh vs Oxy-AZD p = 0.1425; Sal-Veh vs Sal-AZD p = 0.2151. (F) Mushroom spine density was unchanged by oxycodone withdrawal and AZD (Oxy by AZD interaction F(1, 49) = 0.1803 p = 0.6729; Oxy F(1, 49) = 2.390 p = 0.1286; AZD F(1, 49) = 1.999 p = 0.1638). Planned contrasts: Oxy-Veh vs Sal-Veh p = 0.2106, Oxy-Veh vs Oxy-AZD p = 0.2367; Sal-Veh vs Sal-AZD p = 0.4444.

To examine potential behavioral consequences of the above-described effects of CA4 disruption and AZD, we next turned to oxycodone self-administration. Self-administration paradigms allow rodents to control the amount of drug they consume, and thus may be the closest model for drug use and drug-seeking in humans [1, 17]. *Car4^+/+^*mice and *Car4^-/-^* mice were implanted with catheters in the jugular vein for oxycodone self-administration (0.25mg/kg/infusion) in response to active lever presses (ALP) on a FR1 schedule. Mice were allowed to self-administer oxycodone for 6 hours per day for 10 consecutive days (**Fig. 7A**). A 30-minute baseline test of cue-induced drug-seeking behavior was assessed on day 11, in which ALPs produced the light and tone cues but no drug infusion. After 30 days of forced abstinence (experimental day 42), mice received a single administration of AZD (30 mg/kg, i.p.) or vehicle and 24-hours later again underwent a cue-induced drug-seeking session for comparison to baseline testing. Overall, we found mice of both genotypes similarly acquired the self-administration task (days 1 through 10) and both genotypes developed a similar preference for ALPs versus inactive lever presses (ILPs) (**Fig. 7B**). Additionally, both genotypes received a similar number of oxycodone infusions (**Fig. 7C**). These results suggest that CA4 disruption did not affect oxycodone consumption. After 30 days of abstinence, vehicle-injected *Car4^+/+^* mice maintained the same amount of cued drug seeking relative to the baseline (**Fig. 7D**), suggesting that the desire to obtain oxycodone was sustained. In contrast, the AZD-injected *Car4^+/+^* mice reduced their cued drug seeking by half. This reduction was significant compared to vehicle-injected *Car4^+/+^* mice (**Fig. 7D**). Importantly, on day 43, ALPs in *Car4^-/-^*mice were also reduced by half relative to baseline testing (day 11), and AZD had no additional effects in these mice (**Fig. 7D**). Together, these data suggest that after 30 days of abstinence, AZD and CA4 disruption both suppressed cued oxycodone-seeking behavior and that the effects of AZD depended on CA4.

**Figure 7.**
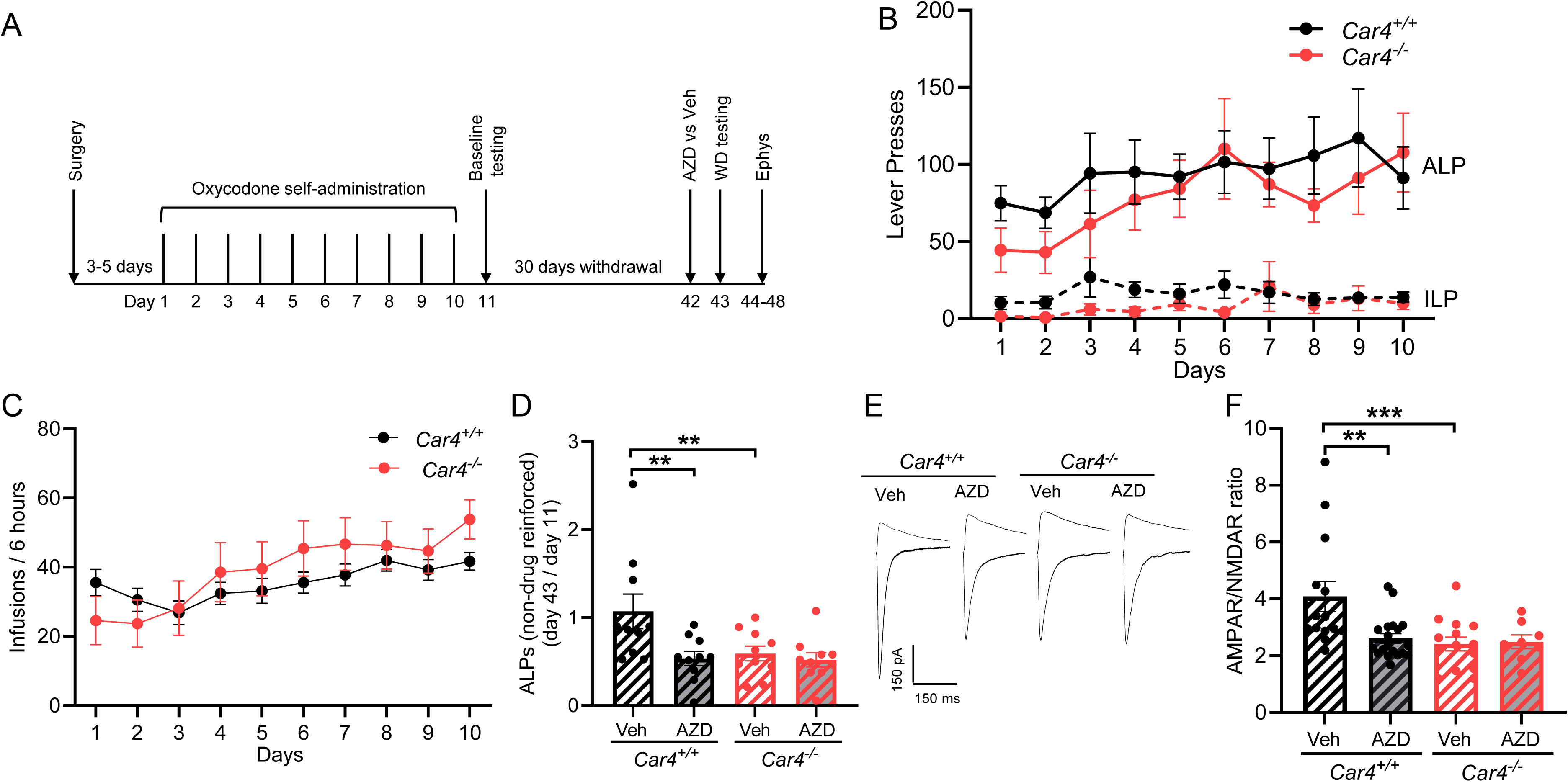
CA4 disruption and inhibition reduced cue-induced drug-seeking behavior following forced abstinence from oxycodone self-administration. (A) Experimental timeline for intravenous oxycodone self-administration and slice electrophysiology. (B) Lever presses (active (ALP) and inactive (ILP) across experimental sessions. There was no effect of genotype (*Car4^+/+^* vs *Car4^-/-^*) on ALP (F(1, 30) = 0.2533, p = 0.6187) or ILP (F(1, 30) = 0.8632, p = 0.3602). n = 10 mice/per group. (C) Infusions across experimental sessions between the genotypes. Interaction F(1, 200) = 1.152 p = 0.3278. There was no effect of genotype (F(1, 20) = 0.4102, p = 0.5291). (D) Cue-induced oxycodone seeking (ALPs) after 30 days of forced abstinence (day 41) normalized to the baseline seeking test 24 hours after the last self-administration session (baseline, day 11). Vehicle treated *Car4^-/-^* mice were reduced ALP compared to vehicle-treated *Car4^+/+^* mice. AZD treatment significantly reduced ALPs in *Car4^+/+^* mice. However, in *Car4^-/-^* mice, AZD treatment did not affect ALPs. Genotype by AZD interaction(F(1, 36) = 3.628 p = 0.0648, treatment(F(1,36) = 6.226 p = 0.0173, genotype (F(1, 36) = 4.161 p= 0.0488. Planned contrast, *Car4^+/+^* Veh vs *Car4^+/+^* AZD **p = 0.0036, *Car4^+/+^* Veh vs *Car4^-/-^* Veh **p = 0.0084, *Car4^+/+^* Veh vs *Car4^-/-^* AZD **p = 0.0028. (E) Representative traces of AMPAR (-70 mV) and NMDAR (+50 mV) from NAcC of *Car4^+/+^*and *Car4^-/-^* mice treated with AZD vs. vehicle and recorded after forced abstinence from oxycodone self-administration and behavioral testing of drug seeking. (F) AZD treatment reversed AMPAR/NMDAR ratio in *Car4^+/+^*mice without affecting *Car4^-/-^* mice. Genotype by AZD interaction(F(1, 51) = 5.126, p = 0.0278. Planned contrast, *Car4^+/+^* Veh vs *Car4^+/+^* AZD **p = 0.0011, *Car4^+/+^* Veh vs *Car4^-/-^* Veh ***p = 0.0006, *Car4^+/+^* Veh vs *Car4^-/-^* AZD **p = 0.0045. n = 8-10 neurons from 3 to 6 mice/group.

We next harvested brain tissue from these same animals on days 44-48, to test whether behavioral effects of AZD and CA4 might be related to the oxycodone withdrawal-induced differences in glutamatergic transmission observed in our prior experiments (**Fig. 3B, D**). Indeed, we found that AZD treatment in *Car4^+/+^* mice significantly attenuated the AMPAR/NMDAR ratio in NAcC-MSNs compared to non-AZD treated *Car4^+/+^*mice (**Fig. 7E, F**). Importantly, AZD treatment did not impact the AMPAR/NMDAR ratio in *Car4^-/-^*mice, and the AMPAR/NMDAR ratio was significantly lower in vehicle-treated *Car4^-/-^*mice compared to vehicle-treated *Car4^+/+^* mice. Together, these results correlate with behavioral changes observed during the drug-seeking test (**Fig. 7D-F**). Further, these observations support the possibility that behavioral effects of AZD and CA4 disruption are closely related to their effects on glutamatergic synapses in NAcC-MSNs.

## Discussion

The results above provide new insights into synaptic adaptations in NAcC MSNs induced by abstinence from opioids, which may contribute to the long-lasting desire for more drug. These results further reveal a novel role for carbonic anhydrase 4 (CA4) in opioid-induced synaptic adaptations and suggest that medications that inhibit CA4, such as AZD, may hold promise for reducing the risk of relapse in people with OUD.

We found that oxycodone withdrawal increased AMPAR/NMDAR ratio and increased CP-AMPARs at glutamatergic synapses in NAcC MSNs in *Car4^+/+^*mice, as reflected by elevated rectification index and NASPM sensitivity. We do not yet fully know the temporal limits of these neuroadaptations, but here changes in AMPAR/NMDAR required more than 24-hrs, emerged within 5 days of abstinence, and persisted for more than 30 days following oxycodone self-administration and were thus relatively long-lasting. The above-described effects of oxycodone occurred with both experimenter-administered and self-administered drug, suggesting that they depended on the drug itself rather than how it was administered. Additionally, the oxycodone-induced increase in AMPAR/NMDAR ratio mapped to D1^+^ MSNs expressing D1R and not to non-D1^+^ MSNs. Because D1^+^ MSNs promote reward- and drug-seeking behaviors [21, 22], this finding aligns with a long-lasting drive for opioids following abstinence from repeated oxycodone exposures.

Our findings that oxycodone effects generalized to other mu receptor agonists, morphine and heroin, are not surprising. More surprising is that our results with these opioids also paralleled, at least in part, those observed previously in NAcC-MSNs following abstinence from cocaine [18, 20]. Despite binding to different targets, opioids and cocaine share some converging molecular effects. For example, exposure to both opioids and cocaine alters dopamine signaling, which has been suggested to contribute to neuroadaptations in glutamate receptor trafficking in NAcC [55]. One effect of cocaine withdrawal that we did not observe with oxycodone withdrawal was an increase in dendritic spine density. We and others have previously observed increases in dendritic spine density in NAc MSNs following abstinence from cocaine [18, 51−53]. However, unlike cocaine, here we found no change in spine density following abstinence from oxycodone, which agrees with another recent study where abstinence from oxycodone did not alter spine density in NAcC but instead reduced spine volume and spine neck diameter [24]. Others have also previously noted differences evoked by morphine versus cocaine on dendritic spines [53]. Thus, although at least some effects of opioids and cocaine on synaptic transmission are shared, effects on dendritic spines seem to diverge.

Importantly, changes in glutamatergic synapse physiology observed following abstinence from opioids were prevented by CA4 disruption. We speculate that these effects of CA4 disruption were due to increased activation of ASIC1A and perhaps its ability to raise intracellular Ca^2+^[18, 20]. We have previously found that disrupting CA4 and pharmacologically inhibiting it with acetazolamide increased ASIC1A-mediated synaptic currents in NAcC MSNs by lowering synaptic pH buffering capacity [18, 20]. Additionally, because loss of ASIC1A in NAcC MSNs increased sensitivity to cocaine- and opioid-induced synaptic rearrangements [20, 24], it seems reasonable to expect that the capability of CA4 disruption to enhance ASIC1A-mediated synaptic currents would produce the opposite effect and reduce sensitivity to drug-evoked synaptic changes. Consistent with the expectation that effects of CA4 disruption depend on ASIC1A, here AZD failed to affect AMPAR/NMDAR ratio in *Asic1a^−/−^* mice.

Acetazolamide blocks CA4, but it also blocks other carbonic anhydrases [31, 42]. Thus, it is possible that the effects of acetazolamide might come from the inhibition of carbonic anhydrases besides CA4. However, if this were the case, we would have expected effects of acetazolamide in *Car4^−/−^* mice as well as in *Car4^+/+^* mice. We saw no effects of acetazolamide in mice lacking CA4, thus strongly suggesting that its ability to reduce oxycodone-evoked synaptic and behavioral responses in wild-type mice is mediated by inhibiting CA4.

Intravenous self-administration remains the gold standard for modeling opioid consumption and seeking behaviors in rodents. Interestingly, CA4 disruption did not affect the acquisition of oxycodone self-administration. Nor did CA4 disruption affect the total number of drug infusions. However, after 30 days of forced abstinence, cue-induced drug-seeking behavior in *Car4^−/−^* mice fell to half of baseline levels. This was in sharp contrast to *Car4^+/+^*mice, which maintained the same amount of active lever presses after 30 days of abstinence compared to baseline. Thus, CA4 disruption weakened cue-induced oxycodone seeking, which was sustained in *Car4^+/+^* mice during an extended period of abstinence. These results echo previous effects of CA4 disruption on drug-seeking and AMPAR/NMDAR ratio following abstinence from cocaine self-administration [18, 20], suggesting that targeting CA4 may reduce seeking behavior for both of these highly addictive drugs. Importantly, administering AZD 24 hrs prior to testing, at a dose used safely in humans, significantly reduced active lever presses in *Car4^+/+^* mice and had no effect in *Car4^−/−^* mice. These results with AZD suggest that inhibiting CA4 acutely can elicit effects similar to chronic CA4 disruption. Because the half-life of acetazolamide in mice is less than an hour [47], the effects of acetazolamide treatment likely lasted beyond its bioavailability. Together, these observations strengthen the possibility that CA4 might be an effective non-opioid therapeutic target for reducing relapse in substance use disorders. Moreover, the AZD results raise the exciting possibility that AZD or similar carbonic anhydrase inhibitors, some already approved for use in humans, might be readily repurposed to help reduce drug-seeking and relapse in OUD and other substance use disorders.

## Acknowledgments

J.A.W. was supported by the National Institute of Drug Abuse (R01DA052953, 5R01DA037216), the Department of Veterans Affairs (Merit Award, IO1BX004440), and the Roy J. Carver Charitable Trust. R.T.L was supported by DA049139 and DA048055. We thank Margaret Fuller for her assistance and the University of Iowa Central Microscopy Research Facility for allowing us to use the Zeiss LSM710 confocal microscope for dendritic spine imaging; this instrument was funded by the NIH (SIG grant, S10RR025439).

## Supplementary Figure Legends

**Figure S1.**
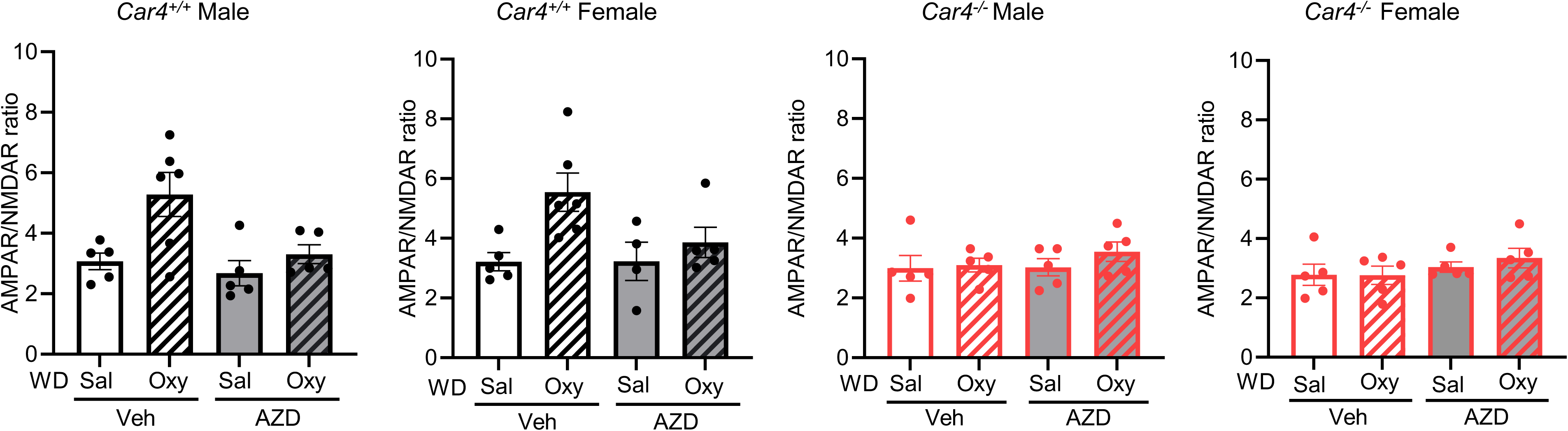
AMPAR/NMDAR ratio results separated by sex, genotype, and AZD treatment. Although there were strong effects of oxycodone withdrawal, *Car4* genotype, and AZD, there were no significant interactions between these factors and sex. Oxycodone by sex: interaction F(1, 18) = 0.011, p = 0.9176; oxy F(1, 18) = 16.03, p = 0.0008; sex F(1, 18) = 0.1292, p = 0.7235. Genotype by sex: interaction F(1, 18) = 0.2851, p = 0.5999; genotype F(1, 18) = 19.41, p = 0.0003; sex F(1, 18) = 0.0045, p = 0.9475. AZD by sex: interaction F(1, 18) = 0.0607, p = 0.8082; AZD F(1, 18) = 9.293, p = 0.0069; sex F(1, 18) = 0.4699, p = 0.5018. n = 5 to 6 neurons from 2 to 3 mice/group.

**Figure S2.**
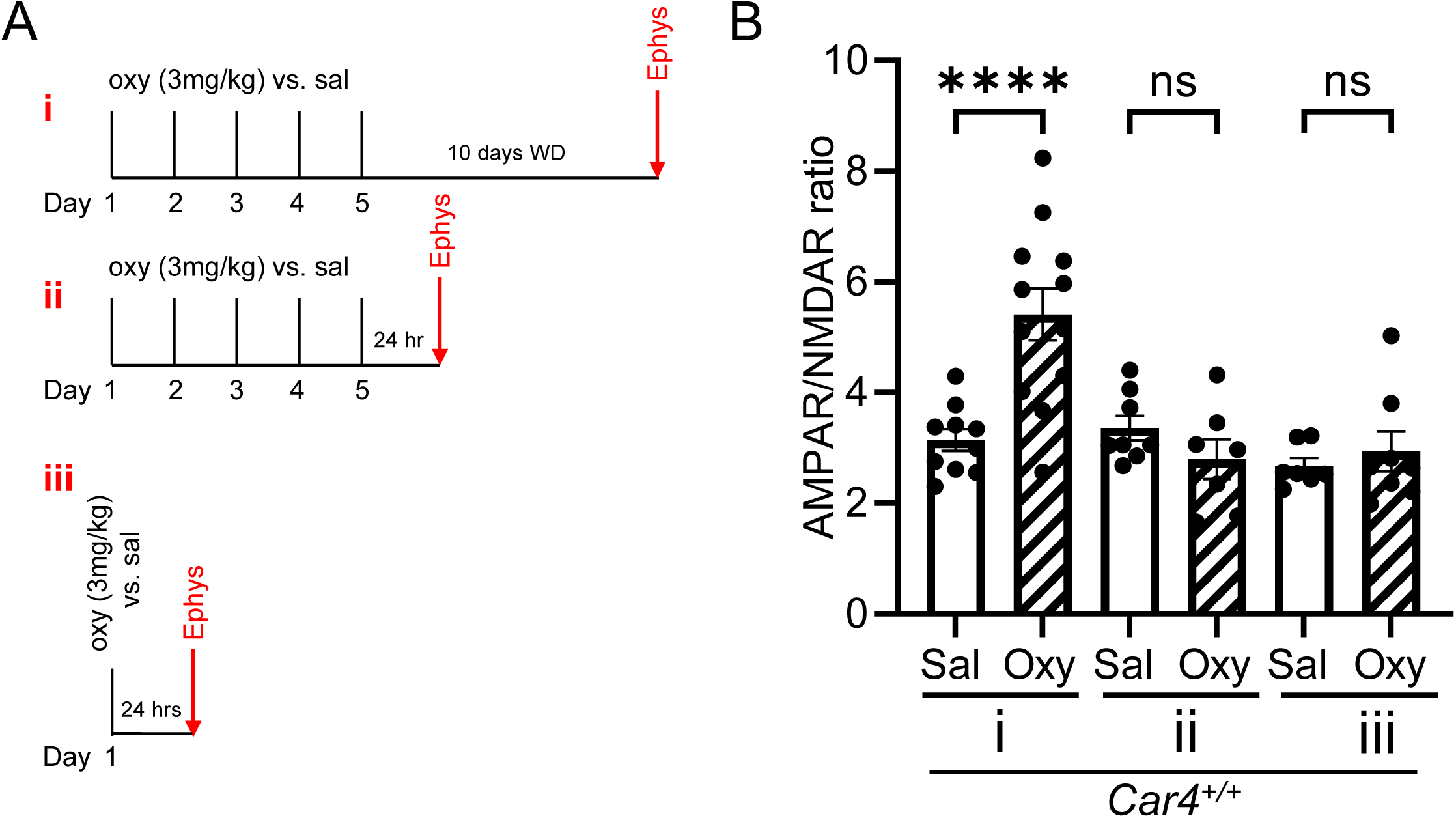
Oxycodone withdrawal-induced increase in AMPAR/NMDAR ratio in NAcC MSNs required an extended abstinence period. (A) Oxycodone (oxy) dosing paradigms administered to *Car4^+/+^* mice to assess effects on AMPAR/NMDAR ratio: (i) 5 injections and 10 days of withdrawal (ii) 5 days of injections and 24 hrs withdrawal (iii) a single injection and 24 hrs withdrawal. (B) AMPAR/NMDAR ratio increased in group (i), but not in groups (ii) or (iii). (i) Sal vs Oxy t(20) = 4.203 ***p = 0.0004, (ii) Sal vs Oxy t(12) = 0.3123 p = 0.7602, and (iii) Sal vs Oxy t(14) = 1.006 p = 0.3318. Students t-test, n = 7-13 neurons from 3 to 4 mice/group.

**Figure S3:**
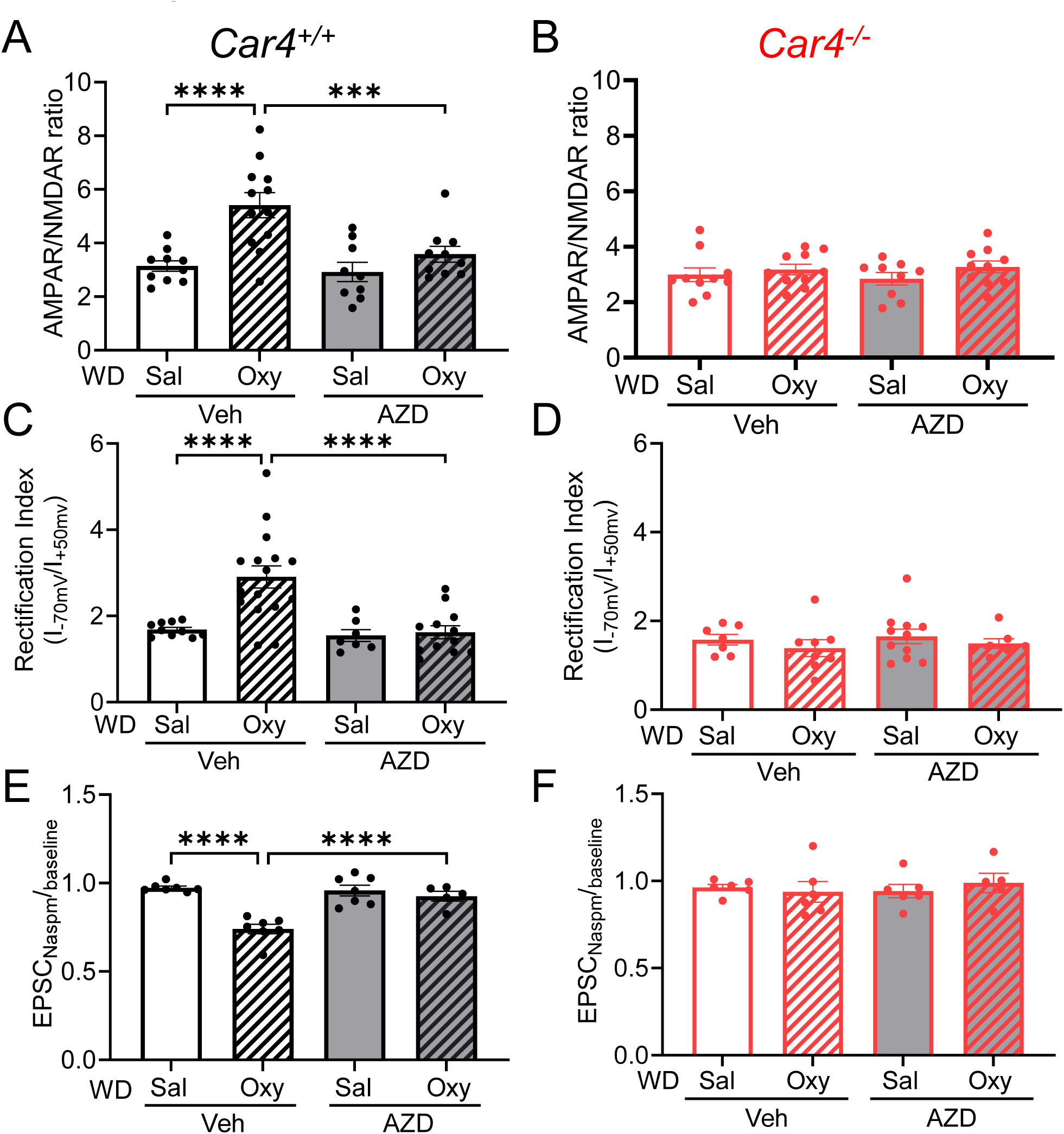
AZD reversed changes at glutamatergic synapse following oxycodone withdrawal in NAcC MSNs in *Car4^+/+^* mice but not in *Car4^−/−^* mice. (A) AZD significantly reduced oxycodone withdrawal-induced increase in AMPAR/NMDAR ratio in *Car4^+/+^*mice (2-way ANOVA, oxycodone by AZD interaction F(1, 37) = 4.966, p = 0.032). Planned contrasts: *Car4^+/+^* Oxy WD-Veh vs Sal WD-Veh ****p < 0.0001, *Car4^+/+^* Oxy WD-Veh vs Oxy WD-AZD ***p = 0.0007. n = 9 to 12 neurons from 3 to 4 mice/group. (B) Oxycodone withdrawal and AZD did not affect AMPAR/NMDAR ratio in *Car4^-/-^* mice (oxycodone by AZD interaction F(1, 35) = 0.2944, p = 0.5908; oxy F(1, 35) = 1.956, p = 0.1708; AZD F(1, 35) = 0.0136, p = 0.9078). Planned contrast; Oxy WD-Veh vs Sal WD-Veh p = 0.5435, Oxy WD-Veh vs Oxy WD-AZD p = 0.7619. n = 9 to 10 neurons from 3 to 4 mice/group (C) AZD treatment attenuated oxycodone withdrawal-induced increase AMPAR rectification in *Car4^+/+^*mice (oxycodone x genotype x AZD interaction F(1, 41) = 6.862, p = 0.0123). Planned contrasts: *Car4^+/+^* Oxy WD-Veh vs Sal WD-Veh ****p < 0.0001, *Car4^+/+^* Oxy WD-Veh vs Oxy WD-AZD ****p < 0.0001. n = 7 to 16 neurons from 3 to 5 mice/group. (D) Oxycodone withdrawal and AZD had no effect on AMPAR/NMDAR ratio in *Car4^-/-^* mice (oxycodone by AZD interaction F(1, 29) = 0.2909, p = 0.5937; oxy F(1, 29) = 0.0007, p = 0.9336; AZD F(1, 29) = 1.16, p = 0.2904). Planned contrast; Oxy WD-Veh vs Sal WD-Veh p = 0.7411, Oxy WD-Veh vs Oxy WD-AZD p = 0.2279. n = 7 to 11 neurons from 3 to 4 mice/group. (E) AZD treatment attenuated oxycodone withdrawal-induced increase in NASPM sensitivity in *Car4^+/+^* mice (Oxy by AZD interaction F(1, 23) = 18.38, p = 0.0003. Planned contrasts: *Car4^+/+^*Oxy WD-Veh vs Sal WD-Veh ****p < 0.0001, *Car4^+/+^* Oxy WD-Veh vs Oxy WD-AZD ****p < 0.0001. n = 6 to 7 neurons from 3 mice/group. (F) Oxycodone withdrawal and AZD had no effect on NASPM sensitivity in *Car4^-/-^* mice (oxycodone by AZD interaction F(1, 19) = 0.6252, p = 0.4389; oxy F(1, 19) = 0.06, p = 0.8092; AZD F(1, 19) = 0.1131, p = 0.7403). Planned contrast; Oxy WD-Veh vs Sal WD-Veh p = 0.6969, Oxy WD-Veh vs Oxy WD-AZD p = 0.4458. n = 5 to 6 neurons from 3 mice/group.

**Figure S4.**
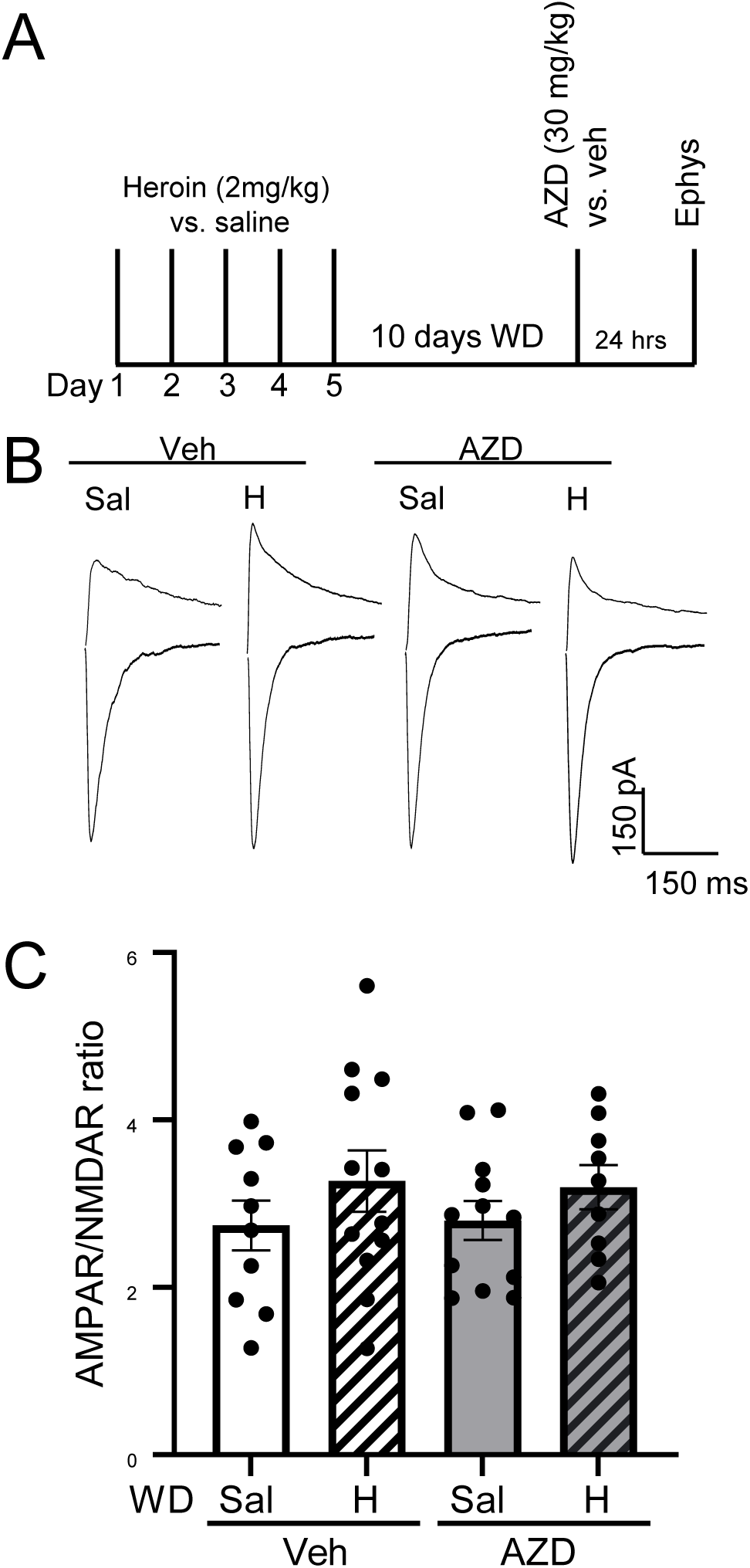
Low-dose heroin (2mg/kg) followed by withdrawal did not affect AMPAR/NMDAR ratio in NAcC MSNs of *Car4^+/+^* mice. (A) Experimental timeline of heroin treatment and electrophysiology (B) Representative traces of AMPAR (-70 mV) and NMDAR (+50 mV) from the saline withdrawal (Sal) and heroin withdrawal (H) treated with vehicle vs. AZD. (C) Abstinence from low-dose heroin did not change the AMPAR/NMDAR ratio and AZD had no effect (Heroin by AZD interaction F(1, 39) = 0.0492 p = 8257, Heroin F(1, 39) = 2.324 p = 0.1355, AZD F(1, 39) = 0.0007 p = 0.9794). Planned contrasts: Sal WD-Veh vs H WD-Veh p = 0.2175, H WD-Veh vs H WD-AZD p = 0.8638, and H WD-Veh vs Sal WD-AZD p = 0.2502. 2-way ANOVA, n = 9-12 neurons from 4 mice/group.

**Figure S5.**
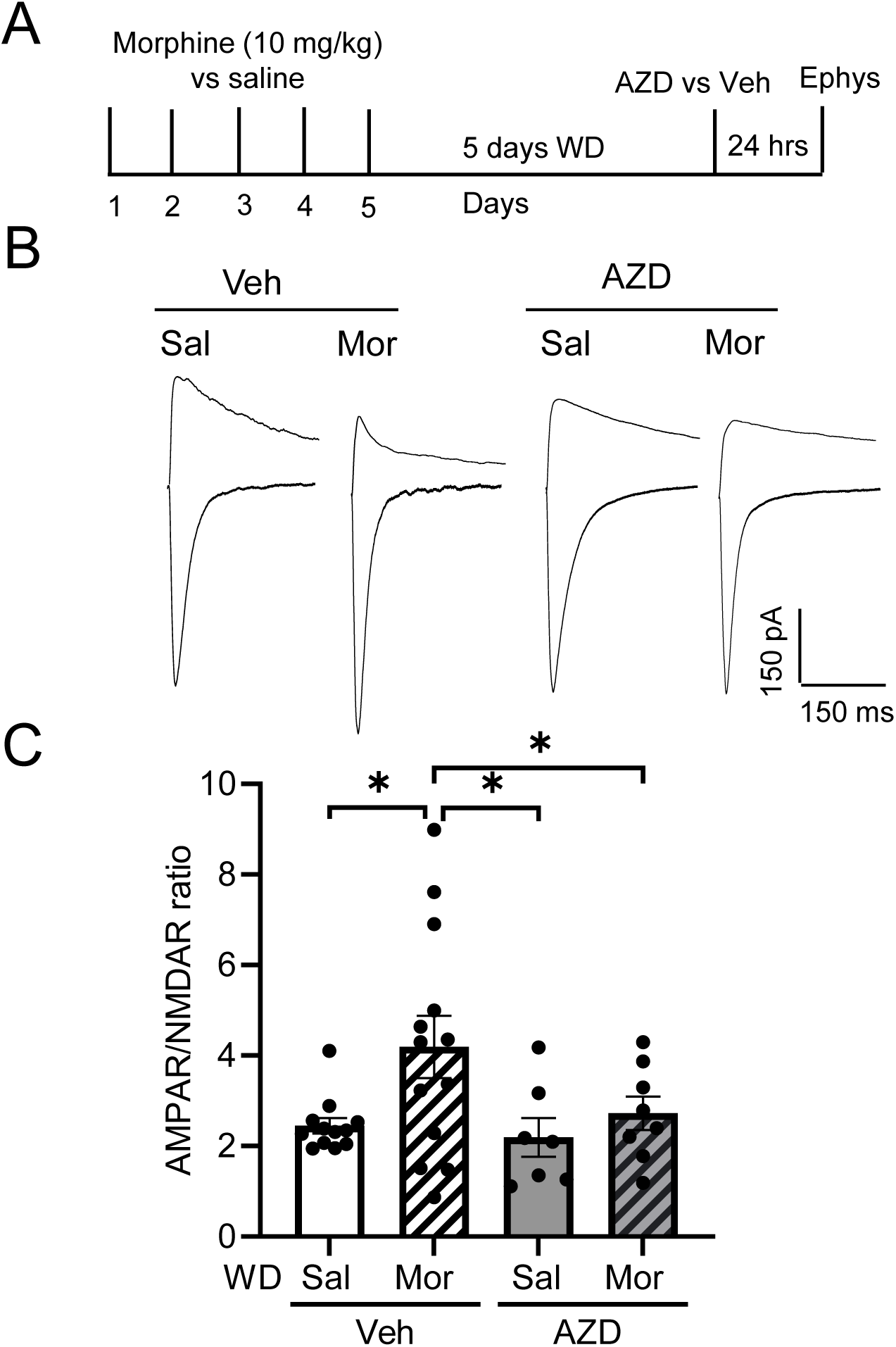
AMPAR/NMDAR ratio was increased in NAcC MSNs of *Car4^+/+^* mice after 5 days of morphine withdrawal, and AZD normalized it to control levels. (A) Experimental paradigm of morphine treatment and electrophysiology (B) Illustrative traces of AMPAR (-70 mV) and NMDAR (+50 mV) from the saline withdrawal (Sal) and morphine withdrawal (Mor) treated with vehicle or AZD (C) AMPAR/NMDAR ratio was increased after 5 days of morphine withdrawal and was normalized by AZD (Morphine by AZD interaction F(1, 36) = 1.327, p = 0.2569; Mor F(1, 36) = 4.659, p = 0.0369; AZD F(1, 36) = 2.697, p = 0.1093). Planned contrasts: Sal WD-Veh vs Mor WD-Veh *p = 0.0102, Mor WD-Veh vs Mor WD-AZD *p = 0.0493, and Sal WD-AZD vs Mor WD-Veh *p = 0.0116. 2-way ANOVA, n = 7-13 neurons from 4 mice/group.

**Figure S6.**
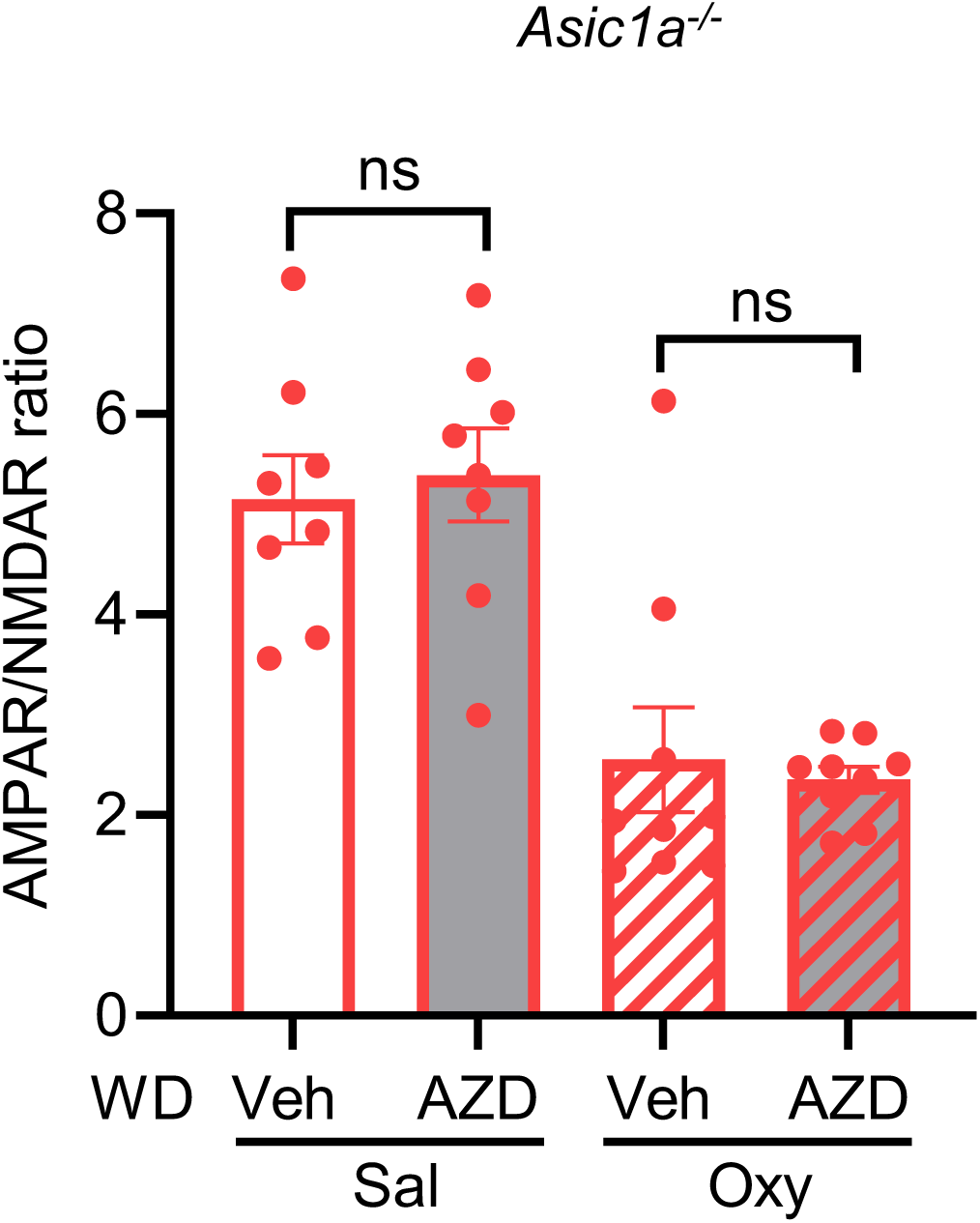
AZD had no effect in drug naïve and oxycodone-withdrawn *Asic1a^-/-^* mice. AZD by oxycodone interaction (F(1, 30) = 0.2836, p =0.5983). Effect of oxycodone (F(1, 30) = 45.97, p < 0.0001. Effect of AZD (F(1,30) = 0.002977, p = 0.9569). Planned contrasts: Sal -Veh vs Sal-AZD p = 0.6895; Oxy -Veh vs Oxy -AZD p = 0.73. n = 8-10 neurons from 4 mice/group.

